# Type V Myosin focuses the polarisome and shapes the tip of yeast cells

**DOI:** 10.1101/2020.07.09.195271

**Authors:** Alexander Dünkler, Marcin Leda, Jan-Michael Kromer, Joachim Neller, Thomas Gronemeyer, Andrew B. Goryachev, Nils Johnsson

**Affiliations:** Institute of Molecular Genetics and Cell Biology, Department of Biology, Ulm University, James-Franck-Ring N27, D-89081 Ulm, Germany

**Keywords:** cell morphology, polar growth, vesicle transport, cortex organization, membrane-less compartments, split-ubiquitin

## Abstract

The polarisome is a cortical proteinaceous micro-compartment that organizes the growth of actin filaments and the fusion of secretory vesicles in yeasts and filamentous fungi. Polarisomes are compact, spot-like structures at the growing tips of their respective cells. The molecular forces that control form and size of this micro-compartment are not known. Here we identify a complex between the polarisome subunit Pea2 and the type V Myosin Myo2 that anchors Myo2 at the cortex of yeast cells. We discovered a point mutation in the cargo-binding domain of Myo2 that impairs the interaction with Pea2 and consequently the formation and focused localization of the polarisome. Cells carrying this mutation grow round instead of elongated buds. Further experiments and biophysical modeling suggest that polarisome nanoparticles use multiple copies of Myo2 and an actin filament polymerizing activity to drive the assembly of the polarisome and sustain its compact shape.

## Introduction

The polar growth of yeasts and other fungal cells is determined by the controlled insertion of plasma membrane and deposition of cell wall material. This material is transported with secretory vesicles on actin cables (Bi and Park, 2012; Jin et al., 2011; Pruyne et al., 2004b; Johnston et al., 1991). The Rho GTPase Cdc42 determines the general polarity of the transport by locally activating proteins that direct the actin cytoskeleton toward the front of the cell (Chiou et al., 2017). During tip growth yeasts and filamentous fungi further concentrate these activities onto a small sector of the cortex. This focus is accomplished by the polarisome, a multi-protein complex that forms a compact protein micro-compartment below the membrane of the growth zone (Sheu et al., 1998; Snyder, 1989; Tcheperegine et al., 2005; Fujiwara et al., 1998; Chenevert et al., 1994). The polarisome combines activities of actin filament nucleation and exocytosis to achieve a spatiotemporal control of vesicle fusion. Composition, structure, and stoichiometry of the polarisome are poorly characterized. Pea2 and Spa2 form the core of this protein assembly and recruit the yeast formin Bni1, the actin nucleator Bud6, and the GAPs for the Rab GTPase Msb3 and Msb4 (Evangelista et al., 1997; Amberg et al., 1997; Arkowitz and Lowe, 1997; Tcheperegine et al., 2005; Moseley and Goode, 2005; Fujiwara et al., 1998; Valtz and Herskowitz, 1996; Sheu et al., 1998). Yeast cells lacking any of these proteins grow round instead of ellipsoid-shaped buds (Tcheperegine et al., 2005; Neller et al., 2015; Sheu et al., 2000). The polarisome also links the kinases of the cell wall integrity pathway to the bud tip to control the inheritance of the cortical ER and to possibly coordinate membrane insertion with cell wall synthesis (Li et al., 2013; van Drogen and Peter, 2002; Sheu et al., 1998; Hruby et al., 2011). Recently, new members and functions of the polarisome were added. Epo1 binds Pea2 and links the cortical endoplasmic reticulum to the cell tip (Chao et al., 2014; Neller et al., 2015). Aip5 binds Spa2 and supports the Bni1-Bud6 complex during actin polymerization (Glomb et al., 2019) (Xie et al., 2019). By assuming a well-delimited compact shape throughout bud formation and expansion, the polarisome shares features with other membrane-less compartments. Like the polarisome, these compartments are thought to regulate cellular pathways by bringing together their major components in one cellular site. Membrane-less compartments with a polar cellular localization have recently been shown to form by nonequilibrium liquid-liquid phase separation regulated by the expenditure of cellular energy (Brangwynne et al., 2009; Hyman and Brangwynne, 2011; Zwicker et al., 2014; Saha et al., 2016). However, other ways to couple the hydrolysis of ATP or GTP to the entropy-reducing formation of polar cellular structures are also known (Goryachev and Pokhilko, 2008; Kirschner and Mitchison, 1986). What are the forces that keep the polarisome’s position, shape and size? Here we discover a general actomyosin-based mechanism that spatially focuses the proteins of the polarisome into a compact micro-compartment.

## Results

### Myosin-V interacts with the polarisome through Pea2

Spa2 and Pea2 form the main structural core of the polarisome complex whose many copies form a sub-micron-sized structure, the polarisome (Figure 1A). The polarisome first emerges at the incipient bud site by a process that appears as a coalescence of small particles of polarisome complexes (polarisome-nanoparticles (PNPs) (movie 1). The polarisome remains associated with the tip of the growing bud and seems to adsorb and to release PNPs throughout the bud growth (Figure 1A, movie 1). Its characteristic focused, spot-like appearance is easily distinguishable from the diffuse spatial distribution of active Cdc42 (Figure 1A). During late G2 the polarisome disintegrates back into PNPs that later in the cell cycle coalesce again at the bud neck to form two parallel rings located on each side of the plane of cell separation (movie 1). One possible explanation for the small size, round shape, and high concentration of proteins within the polarisome is the involvement of energy-consuming mechanisms mediated, for example, by molecular motors. Indeed, the budding yeast type V myosin Myo2, but not the type II myosin Myo1, co-localizes with the polarisome at all stages of bud morphogenesis (Figure 1A) (Schneider et al., 2013; Fang et al., 2010).

**Figure 1.**
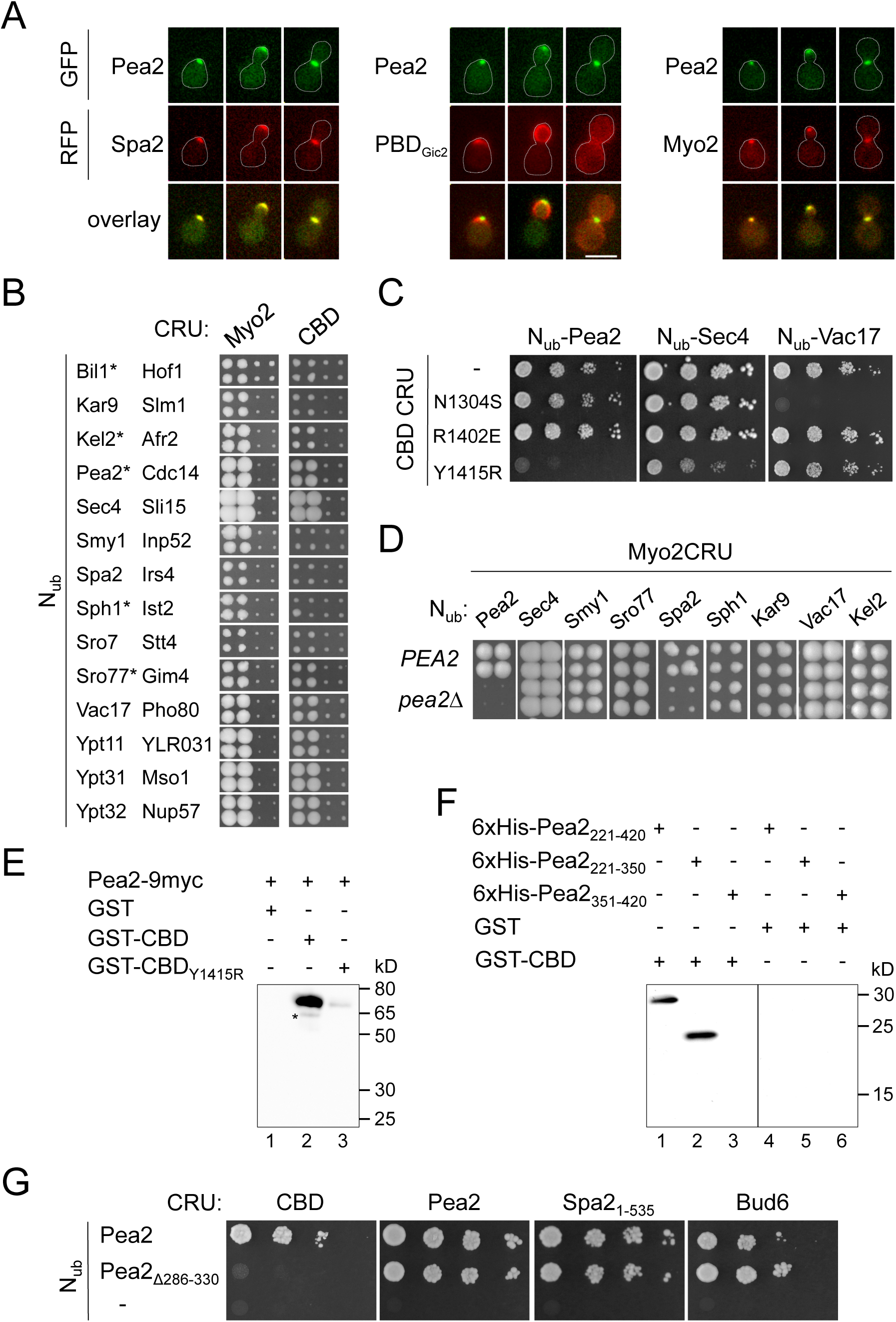
Myo2 binds with its cargo-binding domain (CBD) to the polarisome subunit Pea2. (A) Two-channel microscopy of yeast cells expressing GFP-Pea2 together with Spa2-mCherry (left), Cdc42_GTP_-sensor PBD_Gic2_ (middle), or Myo2-mCherry (right). Shown are images of cells during bud site formation (left column), bud growth (middle column), or mitosis (right column). Scale bar: 5 µm. (B) Cut outs of a Split-Ubiquitin array of diploid yeast cells expressing genomic Myo2CRU (left panel) or a CRU fusion to a plasmid-based cargo binding domain of Myo2 (CBD) (right panel) together with the indicated N_ub_ fusions. Four independent matings were arrayed as quadruplet on media containing 5-FOA. Colony growth indicates interaction between the fusion proteins. Cells co-expressing interacting N_ub_ fusions are displayed on the left row (stars indicate novel binding partners). Cells co-expressing non-interacting N_ub_ fusions are displayed on the right row. Due to the presence of the native Myo2, the interaction signals between CBD CRU and the respective N_ub_ fusions appear always weaker. Complete analysis is shown in Figure S1. (C) Split-Ub assay of cells co-expressing CBD-CRU, or CBD-CRU containing selected point mutations together with the indicated N_ub_ fusions. Cells were grown to OD_600_=1 and 4 µl, or 4 µl of a 10-fold serial dilution spotted on media containing 5-FOA. (D) Split-Ub assay as in (B) but with cells co-expressing Myo2CRU and the indicated N_ub_ fusions in the presence (upper panel, *PEA2*), or absence of *PEA2* (lower panel, *pea2Δ)*. (E) Extracts of yeast cells expressing Pea2-9myc (lane1-3) were incubated with Glutathione-coupled Sepharose beads exposing bacterially expressed GST (lane1), GST-CBD (lane 2) or GST-CBD_Y1415R_ (lane 3). Glutathione eluates were separated by SDS-PAGE and probed with anti-myc antibodies (lane1-3) after Western blotting. The asterisk indicates a degradation product. Ponceau staining of the supernatants after elution and western blot of the yeast extract are shown Figure S2A. (F) Bacterially expressed GST-CBD (lane 1-3) or GST (lane 4-6) coupled to Glutathione Sepharose beads were incubated with bacterially expressed 6xHis-Pea2_221-420_ (lane 1, 4), 6xHis-Pea2_221-350_ (lane 2, 5) or 6xHis-Pea2_351-420_ (lane 3, 6). Glutathione eluates were separated by SDS-PAGE and stained with anti-His antibody after transfer onto nitrocellulose. Inputs of 6xHis-Pea2 fragments, GST-CBD and GST are shown in Figure S2B. (G) Same as in (C) but with haploid cells expressing N_ub_-Pea2, N_ub_-Pea2_Δ286-330_, or N_ub_ as negative control together with CRU fusions to CBD (first panel), Pea2 (second panel), Spa2_1-535_ (third panel), or Bud6 (fourth panel).

Seeking to explore a potential role of Myo2 in shaping the polarisome, we performed a protein-protein interaction screen for binding partners of Myo2. A mating-based Split-Ub array revealed, beside other potential binding partners, a strong interaction signal between Myo2 and Pea2 (Figure 1B, S1; Table S1) (Johnsson and Varshavsky, 1994; Wittke et al., 1999; Dünkler et al., 2012). The interaction could be localized to the cargo-binding domain of Myo2 (Myo2-CBD) (Pashkova et al., 2006)(Figure 1B). Myo2-CBD displays three distinct, partially overlapping binding sites. Site I binds to the Rab GTPases Sec4, Ypt32, Ypt31, Ypt32, the microtubule adaptor Kar9 and the peroxisomal receptor Inn2. Site II binds to the mitochondrial and vacuolar receptors Mmr1 and Vac17, and, finally, site III binds exclusively to the exocyst subunit Sec15 (Jin et al., 2011; Fagarasanu et al., 2009; Eves et al., 2012; Lipatova et al., 2008; Tang et al., 2019). Changing central residues that specifically affect the interaction with known ligands for each site, allowed us to identify binding site I of Myo2 as potential interface for Pea2 (Figure 1C) (Jin et al., 2011). A pull down of Pea2 from yeast extracts with the native or mutated CBD carrying an exchange of the central Y1415 in site I (CBD_Y1415R_) confirmed this finding (Figure 1E, S2A). Split-Ub and pull-down analyses restricted the corresponding interface for Myo2-CBD onto a central element of Pea2 (Pea2_221-350_) (Figure 1F, G). This site is distinct from Pea2’s interface for Spa2, Bud6 or for itself (Figure 1G, S2C). A pull-down of bacterially expressed Pea2 fragments by the GST-coupled CBD of Myo2 confirmed that this interaction is direct and resides between residues 221 and 350 of Pea2 (Figure 1E, S2B).

Our analysis suggests that the loss of Pea2 might abolish the interaction between Myo2 and the subunits of the polarisome but should not affect direct interactions with other partners of Myo2. Accordingly, N_ub_-Spa2 did not interact with Myo2CRU in cells lacking Pea2 whereas the Split-Ub generated interaction signals between Myo2 and Kar9, Sec4, Sro77, Smy1, Kel2, Vac17 or Sph1, a sequence homologue of Spa2, were not visibly altered by the absence of Pea2 (Roemer et al., 1998; Beningo et al., 2000; Korinek et al., 2000; Beach et al., 2000).

### A mutation in the cargo domain of Myo2 disrupts the interaction with Pea2

To obtain mutants of Myo2 that are specifically deficient in their interactions with Pea2, we exchanged side chains of residues surrounding Y1415 on the surface of Myo2-CBD and measured their influence on the interaction with Pea2, the Rab GTPases, or Vac17 (Figures 2A, B, S2D). The Split-Ub binding assays define a hydrophobic patch of three residues in Myo2 that are essential for binding Pea2 and Sec4 (Figure 2A, B). This patch is flanked by a phenylalanine in position 1334 and an arginine in position 1419 that belong to Myo2’s interface to Pea2 but do not contribute to binding the Rab-GTPases or Vac17 (Figure 2B, S2D). F1334, but not R1419, is also part of Myo2’s interface for Kar9 (Eves et al., 2012). To specifically disrupt the Myo2-Pea2 complex we thus focused on position 1419 and replaced the arginine in the genomic *MYO2* by either alanine (*myo2*_*RA*_), or aspartate (*myo2*_*RD*_). As predicted, both mutations selectively impaired the interaction between the full-length Myo2 and Pea2 (Figure 2C). A pull-down of the enriched 6His-tagged Myo2-CBD or Myo2-CBD_RD_ by GST-Pea2_221-350_ confirmed the influence of R1419 on the Myo2-Pea2 interaction *in vitro* (Figure 2D). As *PEA2* is a non-essential gene (Valtz and Herskowitz, 1996), neither of the two mutations in the CBD of Myo2 affected cell growth or survival (movies 1, 2)

**Figure 2.**
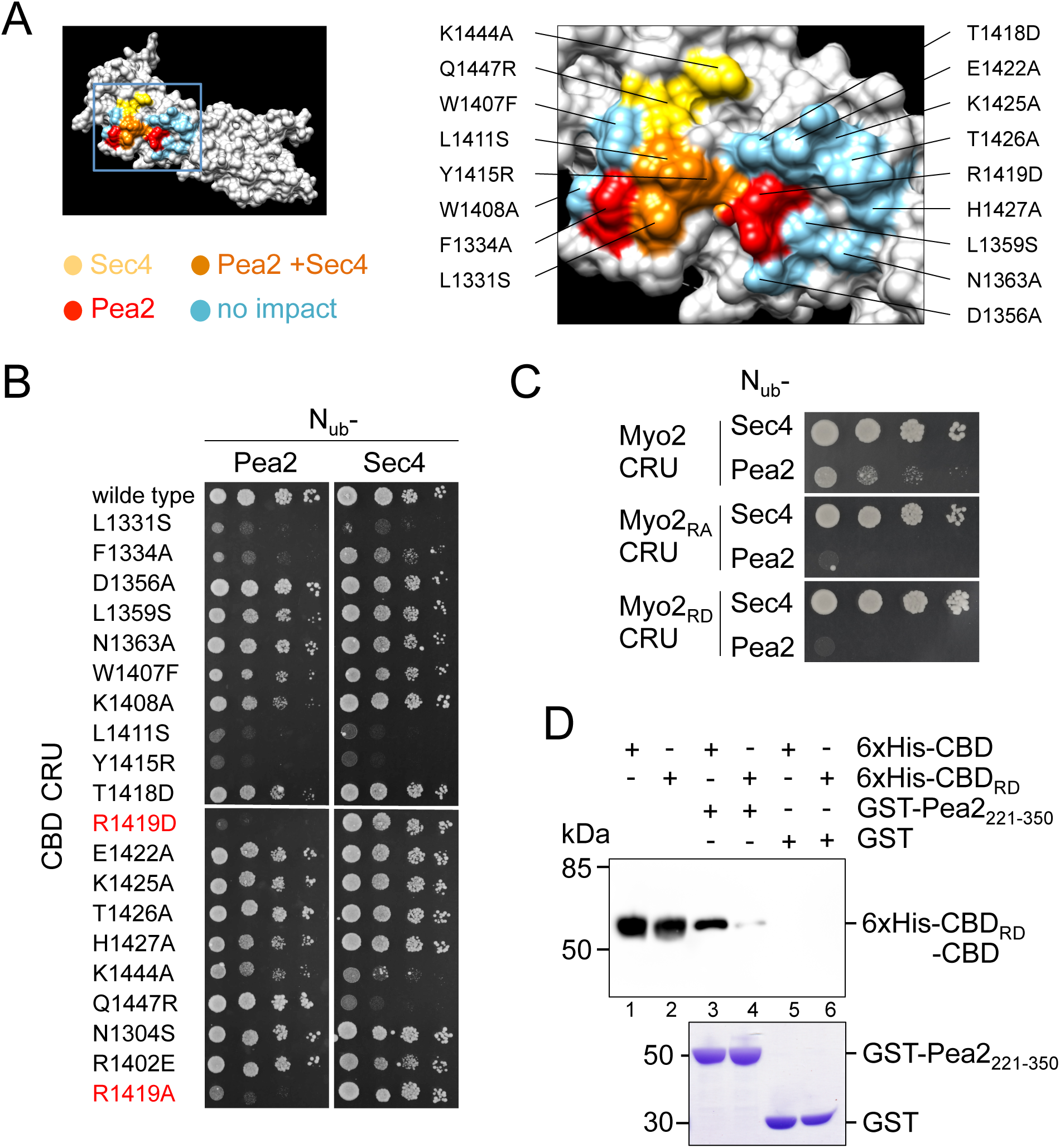
A mutation in the CBD of Myo2 that specifically impairs the interaction with Pea2. (A) Summary of the mutation analysis. Left panel: Structure of the surface of CBD (Pashkova et al., 2006). Residues that exclusively contact Sec4 are colored yellow, those that exclusively contact Pea2 are colored red. Residues contacting both proteins are in orange. Residues with no impact on binding are in blue. Right panel: Blow up of the Myo2 interface showing position and identities of exchanged residues. Color code as in left panel. (B) Dissection of Pea2 binding site on CBD. Yeast cells containing centromeric CBD CRU plasmids with the indicated point mutations and expressing N_ub_-Pea2 or N_ub_-Sec4 were analyzed as in Figure 1C. (C) Split-Ub assay as in (B) but with cells expressing N_ub_-Sec4 or N_ub_-Pea2CRU and fusions to the genomic *MYO2* carrying no (upper panel), the R1419A- (middle panel), or the R1419D exchange (lower panel). (D) Myo2_RD_ affects Pea2 binding *in vitro*. Upper panel: Extracts of *E.coli*-cells expressing 6xHis fusion to CBD (lanes 1, 3, 5) or CBD_RD_ (lanes 2, 4, 6) were purified and incubated with GST-Pea2_221-350_ (lane 3, 4) or GST-coupled beads (lanes 5, 6). Glutathione-eluates were separated by SDS-PAGE, transferred onto nitrocellulose and stained with anti-His antibody. Lower panel: Coomassie staining of the GST fusion proteins after elution from the beads.

### Pea2 anchors Myo2 at the cell cortex to focus growth at the tip

The discovered Pea2-Myo2 interaction could be most readily interpreted as required for the delivery of Pea2 by Myo2 along the actin cables to the bud tip. Indeed, time-lapse fluorescence microscopy shows that Myo2 and Pea2 arrive at the incipient bud site simultaneously (Figure S3, movie S1). Surprisingly, during mitosis, Pea2 arrives at the bud neck shortly before Myo2 (Figure S1). Furthermore, we found that the timing of arrival of Pea2 at both the incipient bud site and the neck are unchanged in cells carrying *myo2*_*RD*_ (Figure S3, movie S1). These observations are inconsistent with the hypothesis that Myo2 transports Pea2 to the polar growth zones.

Careful examination revealed round rather than elliptic bud shapes of the *myo2*_*RD*_- and *myo2*_*RA*_-cells, indicating an even instead of the more pointed bud expansion of wild type cells. The morphology *myo2*_*RD*_- and *myo2*_*RA*_-cells thus demonstrate a striking resemblance to the morphology of *pea2*Δ*-*cells, or cells lacking other core components of the polarisome (Figure 3A) (Sheu et al., 2000; Neller et al., 2015). Accordingly, the GFP fusions to Spa2, Bud6 and Pea2, although still predominantly found at the bud cortex of *myo2*_*RD*_-cells, lacked the pronounced focusing at its center (Figure 3B, movie 2). The cortical fractions of Spa2 and Pea2 were more equally spreaded and the amount of both proteins in the cytosol of mother cells increased (Figure 3B, D, S4B). The comparison between the cellular intensity profiles of Myo2-GFP and its mutant revealed a less prominent tip localization and a higher concentration of Myo2_RD_-GFP in the mother cell (Figure 3B, C, S4B).

**Figure 3.**
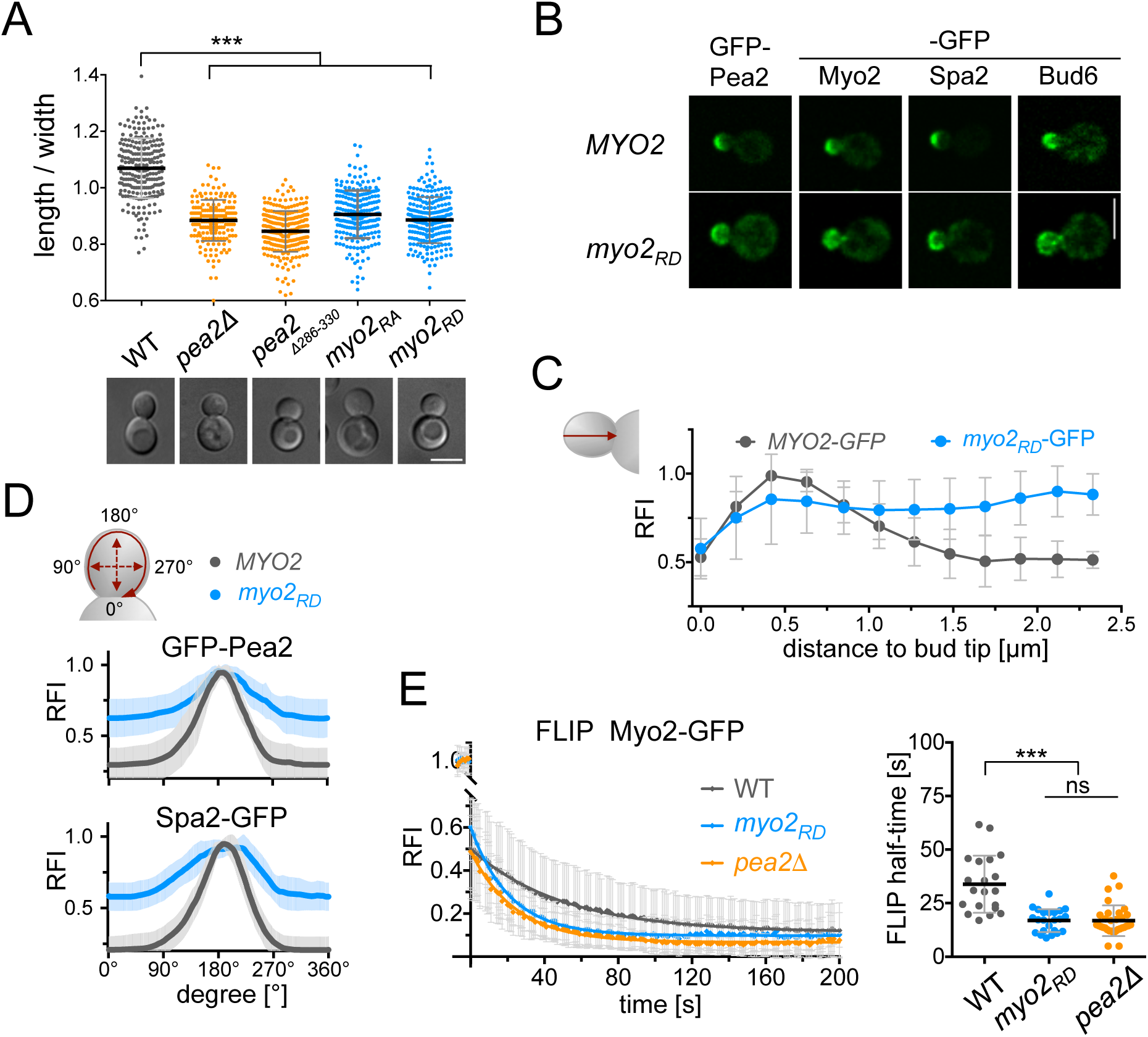
Pea2 recruits Myo2 to the cortex to influence the shape of the daughter cell. (A) Upper panel: Length/width ratios of buds of WT- (mean 1.07; n=244), *pea2Δ-* (0.88 ; n=189), *pea2*_*Δ286-330*_*-* (0.85; n=260), *myo2*_*RA*_*-* (0.91; n=234) and *myo2*_*RD*_*-*cells (0.89; n=247). Error bars indicate SD of the mean. Lower panel: DIC pictures of cells of the corresponding genotypes. Scale bar 5 µm. (B) Representative fluorescence images of small budded WT- or *myo2*_*RD*_-cells expressing GFP-Pea2, Myo2-, Spa2-, or Bud6-GFP. Scale bar 5 µm. (C) Plot profile of the distribution of Myo2-GFP in WT- (grey line, n=12) or *myo2*_*RD*_*-*cells (blue line, n=12). Relative fluorescence intensities (RFI) of single cells were measured along the shortest distance from bud tip to neck (red arrow), and normalized to 1. Error bars indicate SD of the mean. (D) Quantification of GFP-Pea2 and Spa2-GFP fluorescence intensities along the cortex of WT- (grey) and *myo2*_*RD*_- cells (blue). Each oval plot profile (red curved arrow in the upper cartoon) shows the mean of n=12 single normalized profiles. Error bars indicate SD (light grey and blue) of the mean. (E) Disruption of the Pea2-Myo2 complex impairs Myo2’s association with the cortex. Left panel: FLIP analysis of Myo2-GFP in wild type- (n=20), *myo2*_*RD*_- (n=23), and *pea2Δ*-cells (n=32). Photo bleaching of mother cells was done every 5 min and z-stack images were taken every sec. Curves are the fitted mean of single measurements. Right panel: Calculated FLIP half-times of Myo2-GFP in WT- (33.8 s), *myo2*_*RD*_- (19.9 s), and *pea2Δ*-cells (16.8 s). Error bars SD.

Based on these observations, we hypothesized that the interaction between Myo2 and Pea2 might occur predominantly at the cell cortex and play a role in the spatial focusing of the polarisome during bud growth. To quantitatively probe the impact of the Pea2-Myo2 interaction on the cortical anchorage of both proteins we monitored the loss of their GFP fusions from the bud by continuously photo-bleaching the fluorescent population in the mother cell (FLIP). The FLIP revealed that the speed of depleting Myo2-GFP (as measured by t_1/2FLIP_) has roughly doubled in the absence of *PEA2* (Figure 3E; t_1/2FLIP Myo2_=34 sec versus t_1/2FLIP Myo2_ in *pea2*Δ= 17 sec) (Donovan and Bretscher, 2012). Accordingly the t_1/2FLIP_ of Myo2_RD_ was close to the t_1/2FLIP_ of the wild type protein in *pea2*Δ-cells (t_1/2FLIP Myo2RD_= 20 sec versus t_1/2FLIP Myo2_ in *pea2*Δ= 17 sec). At the same time, GFP-Pea2 was more tightly associated with the cortex than Myo2 (t_1/2FLIP Pea2_=72 sec versus t_1/2FLIP Myo2_= 34 sec) and its t_1/2FLIP_ was not affected by the R1419D mutation in Myo2 (Figure S3A; t_1/2FLIP Pea2_ in *myo2*_*RD*_-cells =71 sec). We conclude that Pea2 retains Myo2 at the cortex of the bud. The complex is required for the focused delivery of secretory vesicles to the bud tip.

### The Myo2-Pea2 interaction is sufficiently strong to enable polarisome focusing

Our hypothesis that Myo2-Pea2 interactions contribute to spatial focusing of the polarisome implies that the Pea2-anchored Myo2 drags PNPs on actin cables along the cell cortex. To enable such a motion, binding of Pea2 should activate Myo2 and should be sufficiently strong and persistent. Furthermore, Myo2 bound to Pea2 should retain the full ability to walk along actin cables. To test our hypothesis, we resorted to a Pex3-Pea2 fusion construct that displayed Pea2 on the surface of peroxisomes (Pea2-peroxisomes) (Figure 4A) (Luo et al., 2014; Glomb et al., 2020). If our three assumptions are correct, the cellular localization of Pea2-peroxisomes should be altered by a Myo2-dependent transport along the actin cables. Indeed, wild type cells normally contain on average seven peroxisomes that are uniformly distributed between mother cell and the bud (Figure 4A, B). In contrast, Pea2-peroxisomes were concentrated at one or two spots, typically just below the bud tip (Figure 4A, B, C) Introducing the *myo2*_*RD*_ allele into these cells increased the number of the visually distinct peroxisomes and partially restored their spatially homogenous cellular localization (Figure 4A, B, C). We conclude that Pea2-peroxisomes recruit Myo2 in its active conformation to pull the peroxisomes on actin cables towards the tip of the bud. Thus, the Myo2-Pea2 interaction is sufficiently strong and persistent to also provide force onto PNPs at the cortex of the cell.

**Figure 4.**
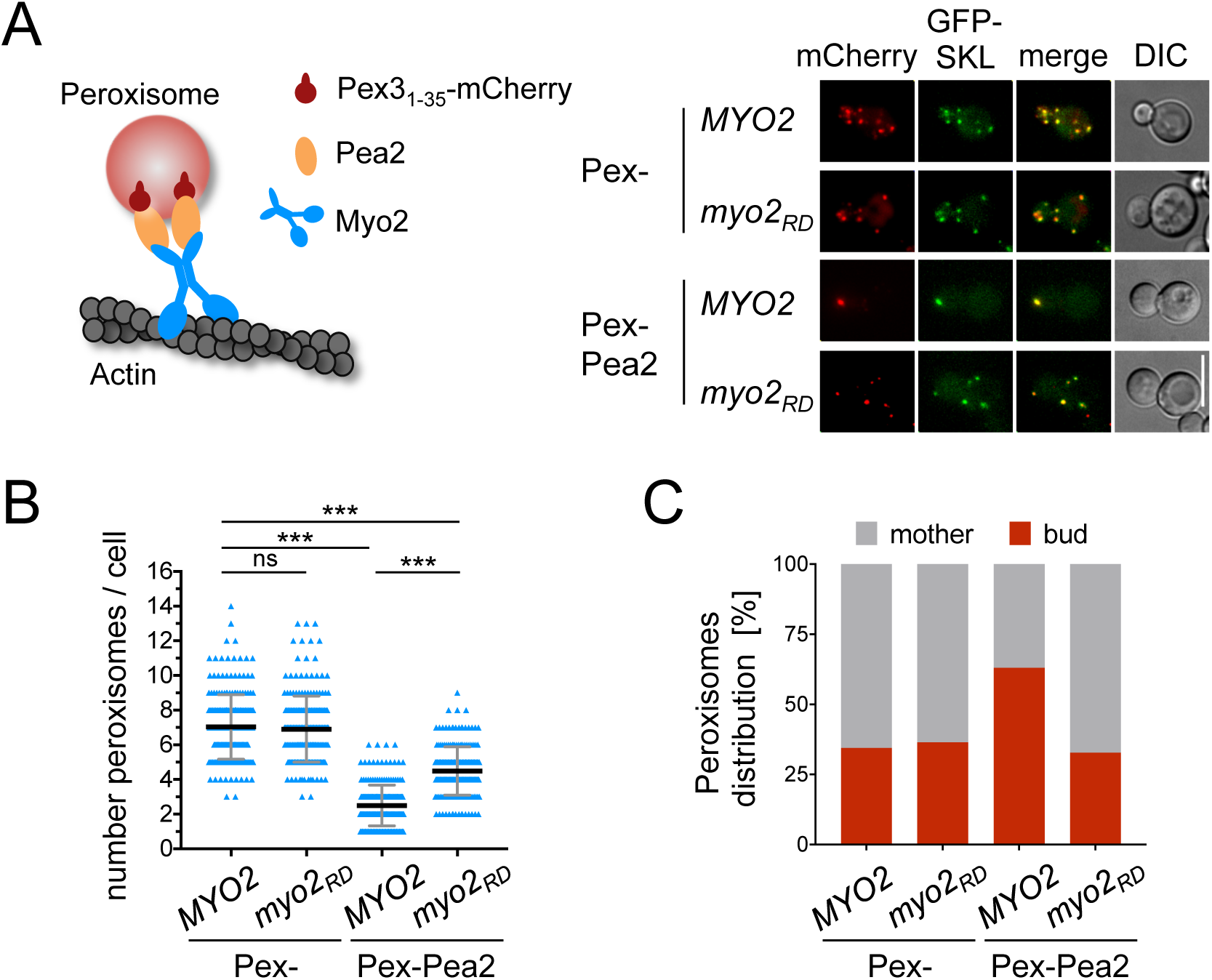
The Pea2-Myo2 complex can walk on actin cables. (A) Left panel: Cartoon of the experimental set up. Pea2 was fused to a fragment of Pex3 and thus coupled to peroxisomes (Pex3_1-35_-mCherry-Pea2 (Pex-Pea2)). Right panel: WT cells or *myo2*_*RD*_-cells co-expressing Pex- or Pex-Pea2 together with the peroxisomal import substrate GFP-SKL were analyzed by two channel fluorescence microscopy. (B) Number of mCherry-stained peroxisomes were quantified in WT- or *myo2*_*RD*_-cells expressing either Pex (n_WT_=228; n_*myo2RD*_=215), or Pex-Pea2 (n_WT_=214; n_*myo2RD*_=215). (C) The distribution of mCherry-stained peroxisomes between mother and daughter cells were determined in WT or *myo2*_*RD*_ cells expressing either Pex (n_WT_=228; n_*myo2RD*_=215), or Pex-Pea2 (n_WT_=214; n_*myo2RD*_=215).

### A biophysical model explains polarisome focusing

To examine under which conditions the interaction between Pea2 and Myo2 could enable spatial focusing of the polarisome, we developed a simple biophysical model. We assumed that PNPs initially occupy an extended cortical domain with the dimensions larger than that of a typical polarisome. We then asked whether the interaction of Pea2-bound Myo2 with dynamic actin cables can result in self-organization of a single, stable polarisome of a submicron size. Following Amberg (Amberg, 1998), and Pruyne and Bretcher (Pruyne and Bretscher, 2000), we assumed that the cables run close to the cortex for ∼1 µm until they diverge from the cortex into the cytoplasm (Figure 5A). In our stochastic model, the PNPs comprise distinct particles of <100 nm in diameter. PNPs include several copies of Pea2-bound Myo2 motors that can walk along a nearby actin cable (Figure 5A, inset). PNPs are also expected to contain formin Bni1 and the nucleation promoting factor Bud6 and, thus, can catalyze the formation of actin filaments and anchor them on their surface. We assume that PNPs either slowly move along the stationary cables, when bound to them, or randomly diffuse on the cortex, when free.

**Figure 5.**
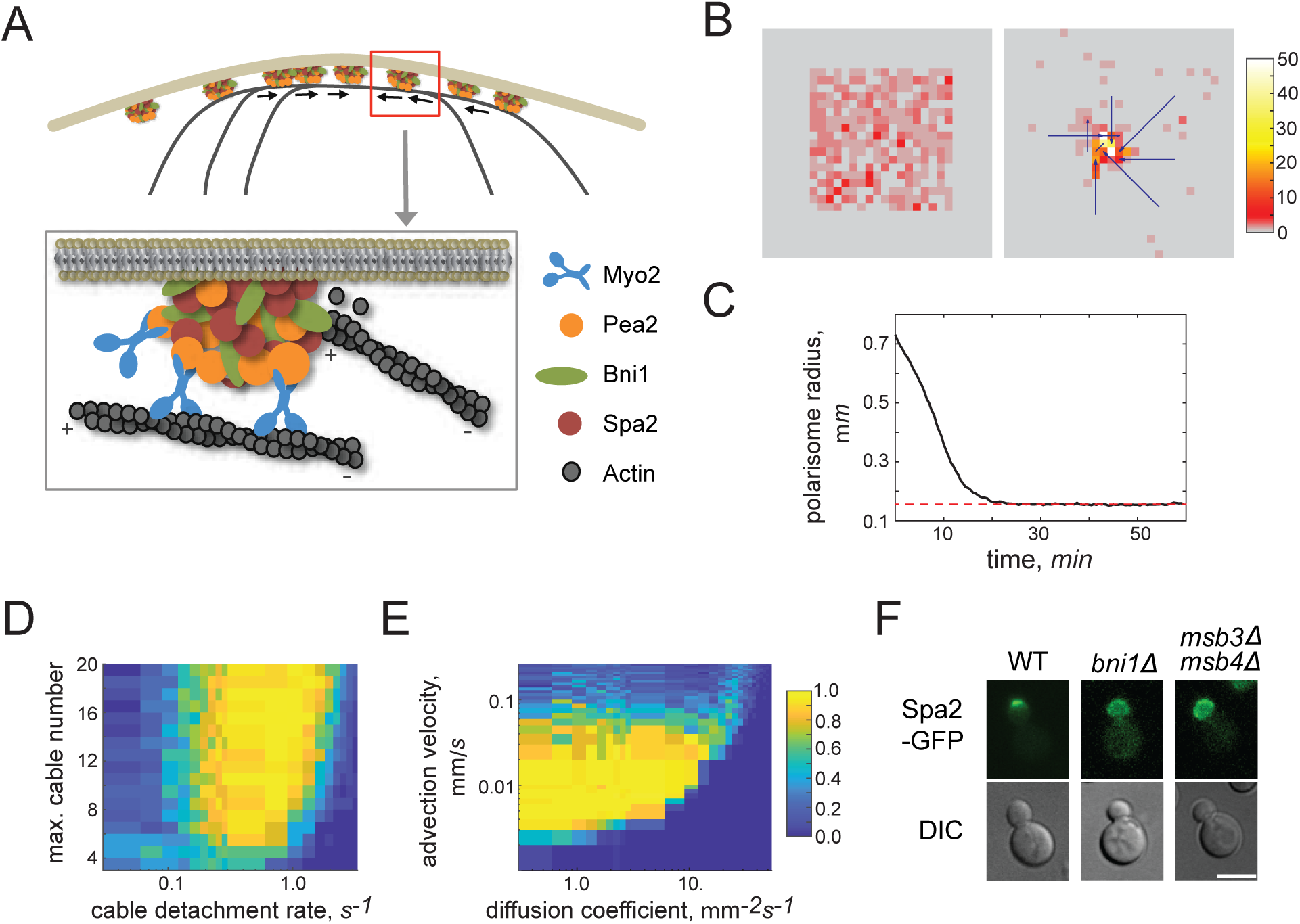
Biophysical model of polarisome focusing. (A) Cortical polarisome nanoparticles move along actin cables via Myo2 motors. (B) Model simulation shows transition of a random initial condition with 0-5 particles per mesh node (left) to the single focused polarisome (right). Actin cables are shown by arrows. Color encodes number of particles per node as indicated by the color bar. Entire simulation is shown in movie 3 (C) Time course of the calculated radius of the polarisome particle cluster. (D,E) Probability to form a single stable polarisome as a function of indicated model parameters. Color encodes the probability as indicated by the color bar. (F) Fluorescence microscopy of yeast cells showing the distribution of Spa2-GFP in wild type- (left panel), *bni1*Δ*-* (middle panel), or *msb3*Δ *msb4*Δ*-*strains (right panel).

We hypothesize the existence of only one positive feedback in the model by assuming that the probability to nucleate a cable at a particular cortical locus is proportional to the number of PNPs at this locus (see Methods for details, Table S2). Once nucleated, the cable can either extend in a random direction, which remains fixed once chosen, or detach and cease to exist. No two cables can nucleate from the same cortical locus and the total number of cables that can exist simultaneously is bounded by *N*_max_. The length of a cable in association with the cortex may not exceed *L*_max_. Together with the probabilities of nucleation and detachment, *N*_max_ and *L*_max_ fully determine the behavior of cables in the model.

Stochastic simulations with extensively varied parameters demonstrated four distinct classes of model behavior: 1) no focusing of PNPs, or formation of 2) multiple dynamic foci, 3) or a single rapidly moving polarisome and, finally, 4) a single stable polarisome. Initial and final states of the model typical of class 4 behavior are shown in Figure 5B. The spatial distribution of cables need not be highly ordered to achieve self-organized focusing of the polarisome (see also movie 3). Time series of the radius of the spatial distribution of polarisome particles (Figure 5C) shows that focusing stops at some characteristic size that remains stable thereafter despite the continuous formation and detachment of dynamic cables.

To analyze the model behavior, we first abrogated the positive feedback by assuming that the cable nucleation probability is independent of the local number of polarisome particles. Under this assumption, cables were nucleated at random spatial cortical locations and produced no focusing of PNPs, regardless of the other parameters. We then systematically varied model parameters and automatically classified the simulation outcomes to determine the probability of formation of a single stable polarisome. The increase in the total number of cables always promotes polarisome focusing (Figure 5D). Variation of the cable detachment rate, however, exhibits a clear optimum. This is because exceedingly dynamic cables fail to substantially alter spatial distribution of polarisome particles before they detach, whereas overly stable cables generate multiple polarisome foci. Thus, intermediate cable lifetimes provide both efficient transport and plasticity necessary for polarisome formation. The focusing is maximal when both the particle diffusion coefficient and their velocity along cables are within their optimal ranges (Figure 5E). Taken together, our modeling results show that the interaction between the polarisome-bound Myo2 motors and actin cables can result in a very efficient spatial focusing of polarisome provided that the dynamics of actin cables and mobility of PNPs are optimized by their co-evolution.

### Actin cables and the Pea2-Myo2 complex are required for polarisome focusing

We next set out to test the predictions of our model. Since actin cables are expected to play the crucial role in focusing the polarisome, their abrogation should result in the polarisome defocusing. Deleting formin *BNI1* (*bni1*Δ) impairs formation of the bud tip-localised actin cables (Pruyne et al., 2004a). Indeed, in complete agreement with our predictions, *bni1*Δ cells exhibited a broadened distribution of Spa2-GFP (Figure 5F). As we already demonstrated that targeted disruption of the Pea2-Myo2 interaction results in polarisome defocusing, we sought an independent approach to test the role of Myo2. Our results showed that Sec4 and Pea2 might compete for binding to Myo2 since they share a common binding interface. Promoting the Sec4-Myo2 interaction should then reduce the Pea2-Myo2 interaction and, therefore, decrease polarisome focusing. Since Myo2 interacts predominantly with Sec4-GTP we deleted the polarisome-localized Sec4 GAPs Msb3 and Msb4 that convert Sec4 into inactive, GDP-bound form (Jin et al., 2011; Gao et al., 2003). This manipulation was already shown to increase the stability of the Myo2-Sec4 complex in cells (Donovan and Bretscher, 2015; Donovan and Bretscher, 2012). Strikingly, the *msb3*Δ *msb4*Δ cells showed broadening of the polarisome identical to that of both the *bni1*Δ and *myo2*_*RD*_ strains (Figure 5F).

### Pea2 recruits Myo2 to the site of cell fusion during mating

Surprisingly, we did not find any effect of *myo2*_*RD*_ on the spatial focusing and size of the polarisome from its emergence at the insipient bud site until the small bud stage (movie 2). This suggests that, at these stages of morphogenesis, the polarisome size is not determined by the interaction between Myo2 and Pea2 and may instead depend on distinct biophysical mechanisms. We thus sought out other morphogenetic scenarios under which formation and size of the polarisome is affected by the Myo2-Pea2 interaction.

Polarisome formation can also be studied during mating of yeast cells (Gehrung and Snyder, 1990). The advantage of studying mating is that the behavior of wild type cells and cells with mutated polarisomes can be compared within the same mating experiment. Here, neighboring a- and α-cells grow mating projections towards each other until both cells get into a physical contact and fuse to form a diploid cell (Figure 6A) (Martin, 2019). The polarisomes form at the prospective mating projections of both cells and stay at their tips during growth and cell fusion (Figure 6A) (Valtz and Herskowitz, 1996; Sheu et al., 2000; Gehrung and Snyder, 1990). GFP-Pea2 and Myo2-GFP accumulated with similar kinetics at the mating projection of wild type cells before post-Golgi vesicles started to expand the mating tip (Figure 6A). When mated with unlabeled wild type cells, Myo2_RD_-GFP appeared considerably later at the prospective mating projection than Pea2-GFP (Figure 6A). The arrival of M*yo2*_RD_-GFP coincided with the arrival of secretory vesicles at the onset of tip expansion (Figure 6A). Subsequent vesicle accumulation and the rate of tip growth were indistinguishable between wild type and mutant cells (Figure 6A).

**Figure 6.**
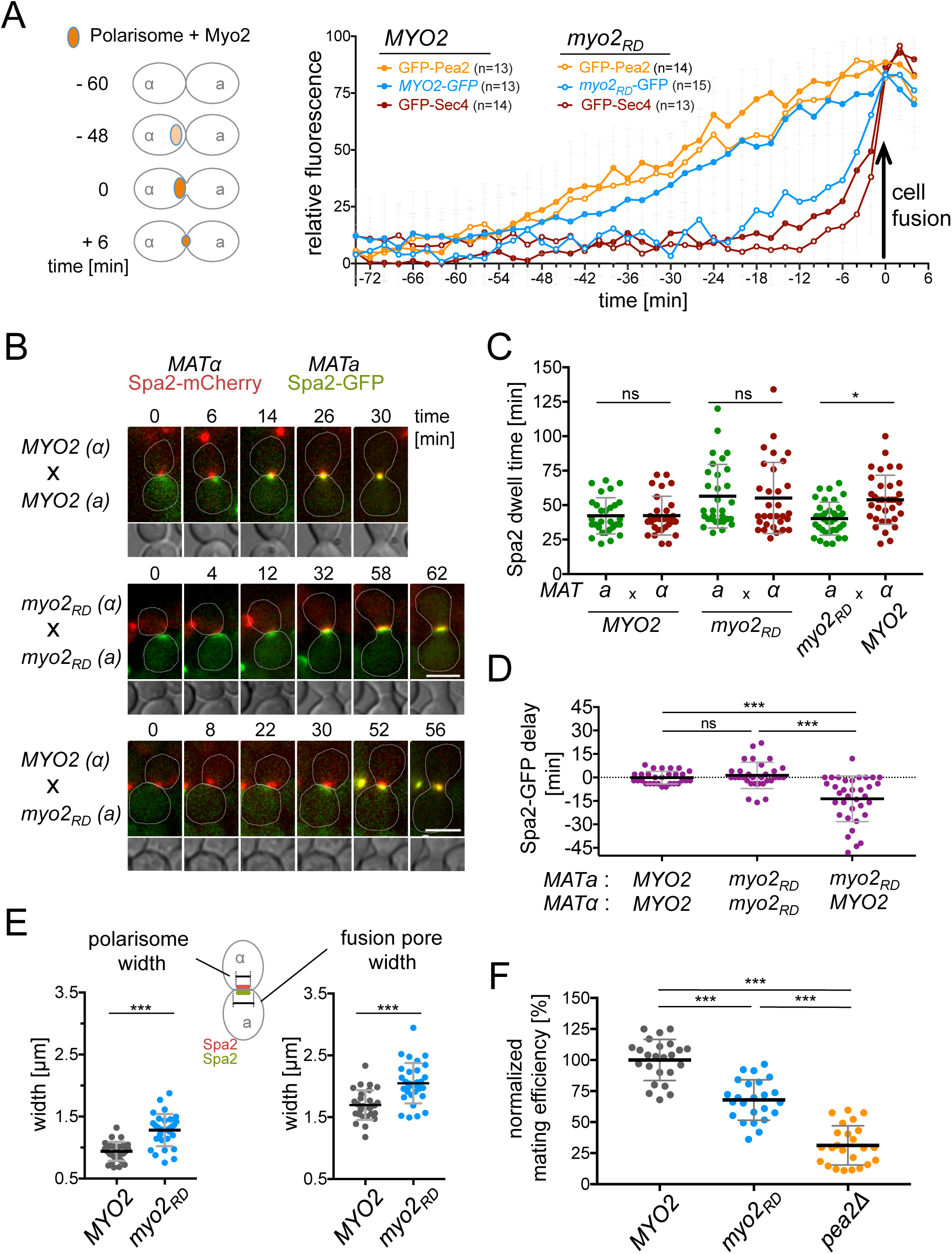
Myo2 stimulates polarisome formation during mating. (A) Left panel: Cartoon of experimental setup. Right panel: *MATa*-cells, or *MATa*-*myo2*_*RD*_-cells expressing GFP-fusion to Pea2 (orange), or Myo2 (not shown), or Sec4 (not shown), were mated with WT *MATα-*cells and the fluorescence intensity was recorded every 2 min. The time of cell fusion was set to t= 0. Right panel: Fluorescence intensity profiles during mating of *MATa*-cells, or *MATa*-*myo2*_*RD*_- cells expressing Myo2-GFP (n_WT_=13; n_*myo2RD*_=15), GFP-Pea2 (n_WT_=13; n_*myo2RD*_=14), or GFP-Sec4 (n_WT_=14; n_*myo2RD*_=13). The highest intensity value of each single curve for each GFP fusion was normalized to 100 to then calculate the mean of the ensemble. Error bars SD. (B) Time resolved fluorescence microscopy (every 2min) of Spa2-GFP-expressing *MATa*- and Spa2-mCherry-expressing *MATα*-cells during mating and cell fusion. The first appearance of Spa2 at the prospective mating projection differed between the genotypes of the mated cells. Upper panel: WT-WT. Middle panel: *myo2*_*RD*_-*myo2*_*RD*_. Lower panel: *myo2*_*RD*_-WT. Each time series starts with the first visible Spa2 signal, the DIC image of the contact sites are shown below. Scale bar 5 µm. (C) Quantification of the data from (B). Comparison of the dwell times of the Spa2-GFP/Spa2-mCherry fluorescence at the mating projections from first appearance to cell fusion: a-WT(Spa2-GFP) X α-WT(Spa2-mCherry): 42.3 min/42.3 min (n=29). a-*myo2*_*RD*_ (Spa2-GFP) X α*-myo2*_*RD*_ (Spa2-mCherry): 56.5 min/55.1 min (n=31). a-*myo2*_*RD*_ (Spa2-GFP) X α-WT (mCherry): 40.3 min/53.9 min (n=35). (D) Quantification of the data from (B). Difference between the first appearance of Spa2-GFP and of Spa2-mCherry measured for each mating experiment of the different genotype combinations. Error bars SD. (E) Left panel: Signal width of Spa2 fluorescence at the fusion site. Right panel: total width of the fusion pores calculated from DIC pictures. Values were derived from matings of WT X WT-cells and *myo2*_*RD*_ X *myo2*_*RD*_-cells, 2 min before fusion. Error bars SD. (F) Efficiency of mating: *MATa* and *MATα* strains (WT X WT; *myo2*_*RD*_ X *myo2*_*RD*_; *pea2Δ* X pea2Δ) were mated for 4 h at 30°C and plated on solid media selecting for diploid zygotes. Colony numbers of diploids from WT-matings were set to 100% efficiency. Error bars SD.

To directly address the influence of Myo2 on the assembly of the polarisome, we followed polarisome formation by comparing the appearance of Spa2-GFP and Spa2-mCherry at the mating projections of interacting a*-* and α-cells (Figure 6B). During mating between wild type cells of opposite mating types, and between *myo2*_*RD*_-cells of opposite mating types, Spa2 appeared in both cell types at roughly the same time (Figure 6B, C, D, movie S3). However, during mating between wild type- and *myo2*_*RD*_-cells, the appearance of the Spa2 spot was delayed by approximately 14 minutes in *myo2*_*RD*_-cells and formed opposite to an already existing polarisome of the wild type cells (Figure 6B, C, D, movie S4). When inspecting mating between a- and alpha *myo2*_*RD*_-cells we not only observed a larger lag between polarisome formation and cell fusion but also broader Spa2-GFP/mCherry-stained regions of the cortex at the fusion zones and wider pores between the fused cells (Figure 6E) (Gammie et al., 1998). Quantitative mating assays showed that the observed defects in polarisome formation correlated with a reduced mating efficiency of *myo2*_*RD*_-cells (Figure 6F).

## Discussion

Taking into account the estimates of secretory vesicle turnover, the target size of vesicle fusion, and the elasticity and the turgor of the cell, one can already calculate the shape and growth of pollen tubes, yeast cells and the hyphae of fungi (Campas and Mahadevan, 2009; Minc et al., 2009; Okada et al., 2013). Finding a quantitative connection between these meso-scale parameters and the molecular properties of the underlying network of proteins are the aim of this and other studies in yeasts (Haupt and Minc, 2018; Abenza et al., 2015).

The width of the polarisome determines the zone of vesicle fusion and thus the shape of the tip (Pruyne et al., 2004b; Sheu et al., 2000; Kohli et al., 2008). Continuous incorporation of new membrane and the subsequent expansion of the cell tip necessitate mechanisms that actively keep the polarisome from spreading or from losing its central position at the cortex. Post-Golgi vesicles are continuously transported by Myo2 on actin cables to the cortex of the growth zone (Schott et al., 2002; Johnston et al., 1991). Myo2 is linked to the incoming vesicles by its interactions with the active form of the Rab GTPase Sec4 and the exocyst component Sec15 (Jin et al., 2011). After tethering at the plasma membrane, Myo2 remains associated with the vesicle for approximately 12-18 sec before GTP hydrolysis relieves Myo2 from the bound Sec4_GTP_ (Donovan and Bretscher, 2015). Decreasing the rate of Sec4_GTP_ hydrolysis extends the residence time of Myo2 at the daughter cell (Donovan and Bretscher, 2015). These observations were taken as evidence that Myo2 stays at the cortex only through its interaction with the tethered vesicle and that its function at the cortex is restricted to support tethering and eventually docking of these vesicles to the plasma membrane. Our data however suggest that Myo2 is additionally anchored by the polarisome subunit Pea2 at the cell tip where it uses its actin dependent force to structure the polarisome and thus the architecture of the vesicle receiving and docking zone. Pea2 will bind to Myo2 only after GTP hydrolysis of Sec4 has cleared the shared binding site and Myo2 is detached from the vesicle (Donovan and Bretscher, 2012). This hand-over mechanism will create a high local concentration of Pea2-Myo2 complexes that pull the associated PNPs towards the center of the micro-compartment where most of the actin filaments are generated. The movement toward the center will concentrate more of the actin nucleation factors Bni1 and Bud6 to the bud tip thus creating a positive feedback loop. Our model explains why the extra stabilization of the Sec4-Myo2 interaction through the loss of the Sec4 GAPs Msb3 and Msb4 dissolves the focused shape of the polarisome and induces the formation of round buds. It also explains why the same phenotypes are observed in cells lacking the polarisome-associated actin nucleation and elongation factor Bni1. In support of our model, Yu et al. observed a Myo2-dependent sliding of actin cables along the cortex of yeast cells (Yu et al., 2011). The required association with the cortex might be provided by the herein discovered connection between Myo2 and the polarisome proteins. The resulting concentration of the plus ends of actin filaments could enforce the pull upon PNPs towards the polarisome center. The proposed mechanism has several biophysical features in common with the Search-Capture-Pull-Release (SCPR) model first proposed by Vavylonis and Pollard for the formation of the fission yeast contractile ring (Pollard and O’Shaughnessy, 2019; Vavylonis et al., 2008). However, in the case of polarisome, the prominence of actin tracks and the particles that use them for congression is reversed. Indeed, the size of polarisome nanoparticles is smaller than that of the *S. pombe* cortical nodes, which are robustly discernible by the fluorescence microscopy, whereas actin cables are thicker and much longer than the single actin fibers that were inferred to connect cytokinetic nodes. Furthermore, unlike the SCPR mechanism that involves contractile type II myosin, polarisome focusing relies on a type V myosin that normally delivers cargo.

The contribution of Myo2 to the de novo formation of the polarisome became evident by studying the mating of yeast cells. In wild type cells, Myo2 and the polarisome arrive together before post-Golgi vesicles accumulate at the prospective mating projection, whereas the delayed arrival of Myo2_RD_ seems to postpone the formation of the polarisome. The extended time between polarisome formation and cell fusion is only observed in cells mating with *myo2*_*RD*_-cells. This effect might be caused by the close communication and synchronization of activities between the two cells during mating. We postulate that the same mechanisms that keep the shape of the mature polarisome might also stimulate the emergence of a new one. This activity might be less important during bud site assembly where formation of the polarity patch is more rigidly defined by landmark proteins and where the instantaneous arrival of vesicles suffice to reinforce the formation of a focused area of tip growth (Lai et al., 2018; Chiou et al., 2017). The when and where of polarisome nucleation might depend on its physical links to Cdc42_GTP_ and/or Bem1 (Evangelista et al., 1997).

The defining droplet-like properties of membrane-less compartments are determined by liquid-liquid phase separation (Banani et al., 2017; Hyman and Brangwynne, 2011). The actomyosin dependence of its shape argues that polarisomes do not belong to this class of bimolecular condensates. However, as many aspects of the exact regulation of size, shape, and composition of the polarisome remain to be explored, it is still possible that the predicted coiled–coil elements in the core components of Spa2, Pea2 and other members of the polarisome drive phase separation during the initial formation of the polarisome scaffold (Sheu et al., 1998; Neller et al., 2015; McSwiggen et al., 2019; Banani et al., 2017; Xie et al., 2019).

## Materials and Methods

### Growth conditions, cultivation of yeast strains and genetic methods

All yeast strains were derivatives of JD47, a segregant from a cross of the strains YPH500 and BBY45, and are listed in Table S3 (Dohmen et al., 1995). Yeast strains were cultivated in SD or YPD media at the indicated temperatures. Media preparation followed standard protocols (Glomb et al., 2020). SD medium for Split-Ub assays contained in addition 1 mg/ml 5-fluoro-orotic acid (5-FOA, Formedium, Hunstanton, Norfolk, UK). Gene deletions and promoter replacements by *P*_*MET17*_ were performed by homologous integration of the cassettes derived by PCR from the plasmids pFA6a-hphNT1, pFA6a-natNT2, pFA6a-kanMX6, pFA6a-CmLEU2 or pYM-N35 (Bähler et al., 1998; Janke et al., 2004). *E.coli* XL1 blue cells were used for plasmid amplification and grown at 37°C in LB medium containing antibiotics. *E.coli* BL21 cells were used for protein production and were grown in LB or SB medium at 37°C.

### Generation of plasmids and yeast strains

Detailed lists of all primers and plasmids from this study are provided in Table S4. Genomic gene fusions were obtained as described (Wittke et al., 1999; Dünkler et al., 2012; Neller et al., 2015; Moreno et al., 2013). In brief, fusions of *GFP, CRU*, or *9xMYC* to *MYO2, SPA2, PEA2*, or *BUD6* were constructed by PCR amplification of the respective C-terminal ORFs without stop codon from genomic DNA. The obtained DNA fragments were cloned via *Eag*I and *Sal*I restriction sites in front of the *CRU*-, *GFP*-, *mCherry*-, or *9xMYC*-module on a pRS303, pRS304 or pRS306 vector (Wittke et al., 1999)The plasmids were linearized using a single restriction site within the C-terminal genomic DNA sequence and transformed into yeast. Successful integration was verified by PCR of single yeast colonies with diagnostic primer combinations using a forward primer annealing in the target ORF but upstream of the linearization site, and a reverse primer annealing in the C-terminal module. Gene deletions were obtained by replacing the ORF through single step homologous recombination with an antibiotic resistance cassette derived by PCR from the plasmids pFA6a-hphNT1, pFA6a-natNT2, pFA6a-kanMX6, pFA6a-CmLEU2 or pYM-N35 (Bähler et al., 1998; Janke et al., 2004).

The insertion of GFP between the *PEA2* upstream sequence and the start codon of *PEA2* was achieved by the Cre/loxP recombination system (Gueldener et al., 2002). A PCR fragment generated from the pGFP-loxP-natCUP sequence was placed in front of the genomic *PEA2* by homologous integration. Subsequent expression of Cre-recombinase (pSH47) deleted the *natNT2-P*_*CUP1*_ sequence and led to the in-frame fusion of GFP to *PEA2*. The *pea2*_*Δ286-330*_ allele was generated by a "delitto perfetto” standard protocol (Storici and Resnick, 2006).

GFP-Sec4 and PBD_Gic2_-RFP were expressed in yeast by genomic integration of plasmids pSec4prom-Sec4-GFP306, or YIp211-GIC2-PBD-1.5tdTomato into the *ura3-52* locus (Glomb et al., 2020). *myo2*_*R1419A*_ (*myo2*_*RA*_), *myo2*_*R1419D*_ (*myo2*_*RD*_) and *MYO2*_*WT*_ alleles were obtained by recombination of PCR fragments generated from plasmids pFA-Myo2-CBD(R1419A)-hph, pFA-Myo2-CBD(R1419D)-hph, or pFA-Myo2-CBD(WT)-hph using a forward primer starting at amino acid1152 and a standard reverse S2 primer annealing to the *MYO2* terminator. The chimeric pP_MET_-Pex3_1-45_-mCherry 306 plasmid was adapted from Luo et al., 2014 (Luo et al., 2014; Glomb et al., 2020). *PEA2* was amplified from genomic DNA and inserted in frame behind the mCherry tag using *Bam*HI/*Sal*I restriction sites. The plasmid was subsequently inserted into the yeast *ura3-52* locus by homologous recombination.

### Mating efficiency

Saturated o/n cultures of wild type JD47 (*MATa*) and ULM53 (*MATα*) cells, or a- and *α*-cells each expressing the alleles *myo2*_*R1419D*_, or *pea2Δ* were diluted to OD_600_= 0.4 and grown for 4 h at 30 °C. Cells were adjusted to an OD_600_ of 1 and equal quantities of JD47 and ULM53 cells mixed and incubated for 4 h at 30 °C in YPD. Cells were diluted 1/20, and 250 µL of each mating mixture were spread on media selecting for diploids. Colonies were counted after 2 d at 30°C.

### *In vivo* Split-Ubiquitin interaction analysis

Large scale Split–Ubiquitin assays were performed as described (Dünkler et al., 2012). A library of 548 different α-strains each expressing a different N_ub_ fusion were mated with a Myo2CRU or a CBD-CRU-expressing a-strain. Diploids were transferred as independent quadruplets on SD media containing 1 mg/ml 5-FOA, and different concentrations of copper sulfate to adjust the expression of the N_ub_ fusions. For small-scale interaction analysis, a- and *α-*strain expressing N_ub_ or C_ub_ fusion constructs were mated, or fusion constructs were co-expressed in haploid strains. Cell were spotted onto SD-FOA medium in four 10-fold serial dilutions starting from OD_600_=1. Growth was recorded at 30°C every day for 2 to 5 days.

Point mutations were introduced into the CBD of *MYO2* by overlap-extension PCR spanning *CBD* (1152-end). PCR products were ligated via *Eag*I/*Sal*I restriction sites between the sequences of the *P*_*MET17*_ promoter and the CRU module on a pRS313 vector (Wittke et al., 1999). The centromeric plasmids were transformed into N_ub_-Pea2, -Sec4, -Ypt11, -Ypt31, -Ypt32 and -Vac17 strains and tested for binding.

### *In vitro* binding assays

#### Yeast cell extracts

Pellets from exponentially grown cultures of Pea2-9myc expressing yeast cells were resuspended in yeast extraction buffer (50 mM HEPES, 150 mM NaCl, 1 mM EDTA) with 1x protease inhibitor cocktail (Roche Diagnostics, Penzberg, Germany), and vortexed together with glass beads (3-fold amount of glass beads and extraction buffer to pellet weight) 10 times for 1 minute, interrupted by short incubations on ice. Extracts were clarified by centrifugation at 16.000 g for 8 min at 4°C and directly used for pull down assays.

#### E.coli extracts

*E. coli* BL21 cells (GE Healthcare, Freiburg, Germany) expressing GST- or 6xHis fusions were grown to an OD_600_=0.8 at 37°C in 2YT or LB (6xHis fusions), or at 18°C overnight in SB medium (6xHis-CBD/ -CBD_RD_). Protein synthesis was induced by the addition of IPTG. 2% ethanol was added in case of 6xHis-CBD- and 6xHis-CBD_RD_ expressing cells. Cell pellets were processed directly or stored -80°C. Pellets were washed once with 1xPBS, resuspended in 1xPBS containing protease inhibitor cocktail (Roche Diagnostics, Penzberg, Germany) and lysed by lysozyme treatment (1 mg/ml, 30 min on ice), followed by sonication with a Bandelin Sonapuls HD 2070 (Reichmann Industrieservice, Hagen, Germany). Extracts were clarified by centrifugation at 40,000 g for 10 min at 4°C.

GST-Pea2_221-350_, -Pea2_221-420_ and Pea2_351-420_ were purified by affinity chromatography on a 5 ml GST-Trap column on an Äkta purifier chromatography system, followed by size exclusion chromatography on a Superdex 200 16/60 column (GE Healthcare) vs. HBSEP buffer (10 mM HEPES pH 7.4, 150 mM NaCl, 3 mM EDTA, 0.05% Tween 20). For pulldown analysis, the protein was buffered into PBS using a NAP5 column (GE Healthcare, Freiburg, Germany).

6xHis-CBD and -CBD_RD_ were lysed in IMAC Buffer A (50 mM KH2PO4 pH8.0, 300 mM NaCl, 20 mM imidazole) by lysozyme treatment and sonication, followed by IMAC purification on an Äkta purifier chromatography system using a 5ml HisTrap Excel column (GE Healthcare, Freiburg, Germany). The column was washed by a linear imidazole gradient (20-70 mM), followed by elution in 200 mM imidazole. Eluted proteins were subsequently transferred into PBS buffer using a PD10 column, concentrated and used for pulldown.

#### Binding assay

All incubation steps were carried out under rotation at 4°C. GST-tagged proteins were immobilized on 100 μl Glutathione**–**Sepharose beads in 1xPBS (GE Healthcare, Freiburg, Germany) directly from *E. coli* extracts. As control, GST alone was immobilized in equal amounts. After incubation for 1 h at 4°C with either yeast extracts or purified proteins, the beads were washed three times with the respective buffer. Bound material was eluted with GST elution buffer (50 mM Tris, 20 mM reduced glutathione) and subjected to SDS-PAGE followed by Coomassie Blue staining and Western Blot analysis using an anti-His antibody (Sigma-Aldrich, Darmstadt, Germany) or anti-MYC antibodies (Hruby et al., 2011).

### Fluorescence microscopy

Microscopic observations were performed with an Axio Observer spinning disc confocal microscope (Zeiss, Göttingen, Germany) equipped with an Evolve512 EMCCD camera (Photometrics, Tucson, USA), a Plan-Apochromat 63X/1.4 Oil DIC objective, and 488 nm and 561 nm diode lasers (Zeiss, Göttingen, Germany). Operations were performed with the ZEN2012 software package (Zeiss). Time-resolved imaging was performed with a DeltaVision system (GE Healthcare, Freiburg, Germany) provided with an Olympus IX71 microscope (Olympus, Münster, Germany), equipped with a Cascade II 512 EMCCD camera (Photometrics, Tucson, USA), a 100xUPlanSApo 100×1.4 Oil ∞/0.17/FN26.5 objective (Olympus, Münster, Germany), a steady state heating chamber and a Photofluor LM-75 halogen lamp (Burlington, VT, USA). Exposure time was adapted to every individual GFP and mCherry labeled protein to reduce bleaching and photo toxicity.

Yeast cultures were grown overnight in SD media, diluted 1:8 in 3-4 ml fresh SD medium and grown for 2 to 6 h at 30°C to mid-log phase. 1 ml cell culture was spun down and resuspended in 50 µl remaining medium. 3 µl of this suspension were transferred onto a microscope slide, covered with a glass coverslip and inspected. For time-lapse analysis, cells were immobilized with a coverslip on custom-designed glass slides containing solid SD medium with 1.8 % agarose. For temporal and spatial analysis of mating events, a- and α-cells were grown in SD to mid-log phase and equal amounts of cells were mixed shortly before agarose slide preparation. Images were obtained with a series of 7 or 5 z-slices and a distance of 0.4 or 0.7 µm between each z-slice.

### Photobleaching experiments

FLIP (Fluorescence Loss In Photobleaching) experiments were performed using a Axio Observer spinning-disc confocal microscope (Zeiss, Göttingen, Germany), equipped with a Zeiss Plan-Apochromat 63Å∼/1.4 oil DIC objective, a 488 nm diode lasers and an UGA-42 photo-manipulation system (Rapp OptoElectronic, Wedel, Germany) (Phair et al., 2004). Images were acquired at room temperature every second with a series of 5 z-slices, each separated by 0.4 µm and an excitation power of 12 to 20% depending on the strain. Six image layers were taken before the ROI (mother cell) was initially bleached with 60 to 80% laser power. 4 to 5 image layers were subsequently acquired and bleaching events continuously repeated each 6 sec until a stable plateau of fluorescence intensity in the daughter cell was reached. Data processing (sum-projection and fluorescence signal quantification) was performed using Fiji 2.0. All data sets were double normalized using Excel. The software Prism 6.0 (GraphPad) was used for the fitting of the double normalized data to a one-phase association curve.

### Quantitative analysis of microscopy data

Microscopy files were analyzed and processed using ImageJ64 1.49 or Fiji 2.0 (US National Institute of Health). Images were acquired as adapted z-series and SUM-projected to one layer. For the quantitative comparison of the mean fluorescent intensities, two regions of interest (ROIs) were determined: firstly the signal of interest (e.g. tip, bud neck), and secondly a region within the cytosol. The mean grey values of both ROIs were quantified and background intensity (cytosol) subtracted. Relative Fluorescence Intensity (RFI) values were obtained by normalizing measured values to 1 or 100.

Fluorescence signal distribution within the bud was obtained by analyzing the images using ImageJ64 plugins "line profile” or “ovale plot profile”. Oval plot profile settings were “Number of points = 360” and “Maximum Intensity”. Fluorescence signal values were normalized to 1 and the average of n=12 single measurements calculated.

### Statistical evaluation

All statistical evaluations were performed using Prism 7 (GraphPad). D’Agostino and Pearson normality tests were used to analyzes the distributions of the data sets. Gaussian distributed data were analyzed by t-test, whereas Mann-Whitney test was used for data that did not pass this criterion. For comparison of more than two data sets one-way ANOVA, or Kruskal-Wallis ANOVA tests were applied.

### Modeling

In our model we assumed that Myo2 and Pea2 create a stable complex, which is associated with the polarisome particles that also contain Bni1 and other polarisome proteins. The particles are confined to the cell cortex and may freely diffuse or move along actin cables. Since the particles interact with the plasma membrane and contain multiple copies of several polarisome proteins, both their myosin-motored advection velocities and the free diffusion coefficients are expected to be considerably smaller than the typical velocity of a single Myo2 motor (∼2 µm/s) and the characteristic diffusion coefficient of a membrane protein (0.01-0.1 µm^2^/s), respectively (Table S2). Simulations are performed on a rectangular 2D domain of size *N* _*x*_ × *N* _*y*_ (where *N*_*x*_ = *N*_*y*_ = 30 and the mesh size *dx* = 0.1*μm*) with the periodic boundary conditions.

Our stochastic model is based on the following assumptions:

1. The probability of cable nucleation at a particular cortical locus (mesh node) is proportional to the number of polarisome particles at this node with the coefficient *k*_*n*_ (Table S2). Upon cable nucleation, one of the eight possible elongation directions (along the mesh dimensions and diagonally) is selected from the uniform random distribution. Cable nucleation may occur only in the bin which is not occupied by any other cable.
2. A cable elongates with the rate *k*_*e*_ until it reaches the length *L*_max_ after which it is assumed that the cable elongates into the cytoplasm, which is not simulated explicitly (Table S2).
3. Cable detachment occurs with the rate *k*_*d*_. Detached cables are assumed to drift into the cytoplasm and all polarisome particles that were attached to it prior to detachment are retained at their respective mesh positions (Table S2).
4. Advection of polarisome particles along cable occurs with the rate *k*_*a*_*n* where *n* is the number of particles at the mesh node (Table S2). We do not explicitly consider particles bound to a cable and free particles as separate species. Instead, we assume that binding to a cable is in the fast equilibrium so that the number of bound particles is equal to *n*_*b*_ = *n K* (1+ *K*) and the number of free particles is equal to *n*_*f*_ = *n* (1+ *K*), where *n*_*b*_ + *n*_*f*_ = *n* and *K* = 0.5 is the binding equilibrium constant. Using this approximation, we compute effective diffusion and advection rates

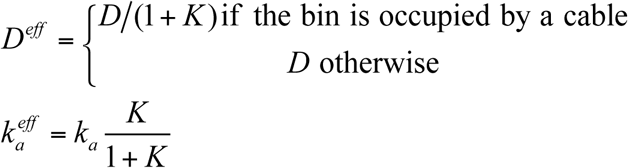

Stochastic simulations are performed using first reaction Gillespie algorithm. At each simulation step tentative times were computed for the following events: 4*N*_*x*_ *N*_*y*_ diffusion events, *N*_*x*_ *N* _*y*_ cable nucleation events, *N*_*c*_(*t*) cable elongation events, *N*_*c*_(*t*) cable detachment events, and 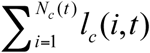 cable advection events. *N*_*c*_(*t*) is the current total number of cables and *l*_*c*_ (*i,t*) is the current i-th cable length. A process with the minimal tentative time is selected and the system state is updated. Time is then advanced by last minimal tentative time. Focal accumulation of polarisome particles was considered stable if its center of mass exhibited only a small-amplitude motion confined within a circular domain with diameter <0.5 um.

Phase diagrams on Figure 5 (D, E) were created as follows. Simulations were run for 60 min of model time. Then temporal averages of the number of particles for the last 1 min for each mesh node were created. The resulting matrix *N*(*i, j*) was then binarized with the threshold equal to 10 (0, if below 10 particles per mesh node, 1 otherwise). If the given stochastic simulation produced a single cluster of ones, then it was assumed that the simulation converged to a single stable polarisome and was scored as 1, otherwise 0. Clustering was performed using MATLAB bwconncomp function. Figure 5 (D, E) presents the probability to converge to a single stable polarisome computed over 30 stochastic realizations for every combination of the shown model parameters.

## Acknowledgement

We thank Nicole Schmid and Steffi Timmermann for assistance during strain construction and protein interaction analysis, and Oliver Glomb for advice during the peroxisome experiments.

## Author contributions

Conceptualization: N.J., A.D., A.B.G; Methodology: A.D., T.G.; Investigation: A.D., M.L., J.-M.K., J.N., T.G.; Writing - original draft: N.J.; Writing - review & editing: N.J., A.D., A.B.G; Supervision: N.J., A.D., A.B.G.; Funding acquisition: N.J., A.B.G.

## Legends to the Figures

**Movie 1**

Time-lapse analysis of wild type yeast cells expressing Spa2-mCherry. Frames were taken every 2 min and cells incubated at 30°C. Scale bar 5 µm.

**Movie 2**

Time-lapse analysis of *myo2*_*RD*_ cells expressing Spa2-mCherry. Frames were taken every 2 min and cells incubated at 30°C. Scale bar 5 µm.

**Movie 3**

Biophysical model of polarisome focusing

## Supplementary Information

### Supplementary Figures

**Figure S1.**
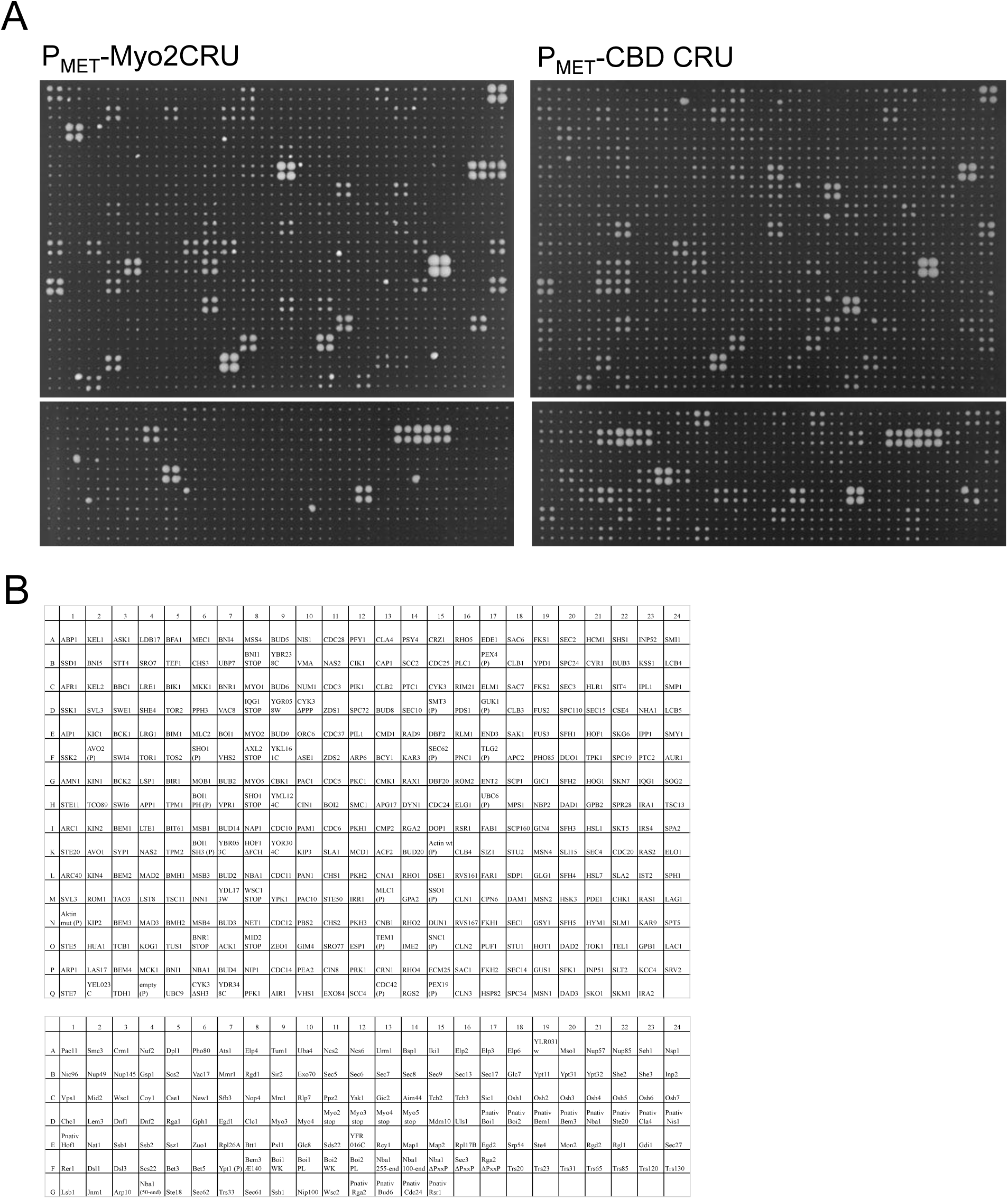
(A) Split-Ub interaction assay of 548 yeast strains co-expressing *P*_*MET17*_-Myo2CRU (left panel) or *P*_*MET17*_*-*CBD CRU (right panel) each with a different N_ub_ fusion protein. In each case cells of four independent matings were spotted as quadruplet on SD medium containing 5-FOA, 150 µM copper sulfate and 70 µM methionine. Shown is the growth of the diploid yeast cells after 3 days at 30°C. Growth indicates protein-protein interaction. (B) Matrix revealing the identities of the respective N_ub_-fusions of (A).

**Figure S2.**
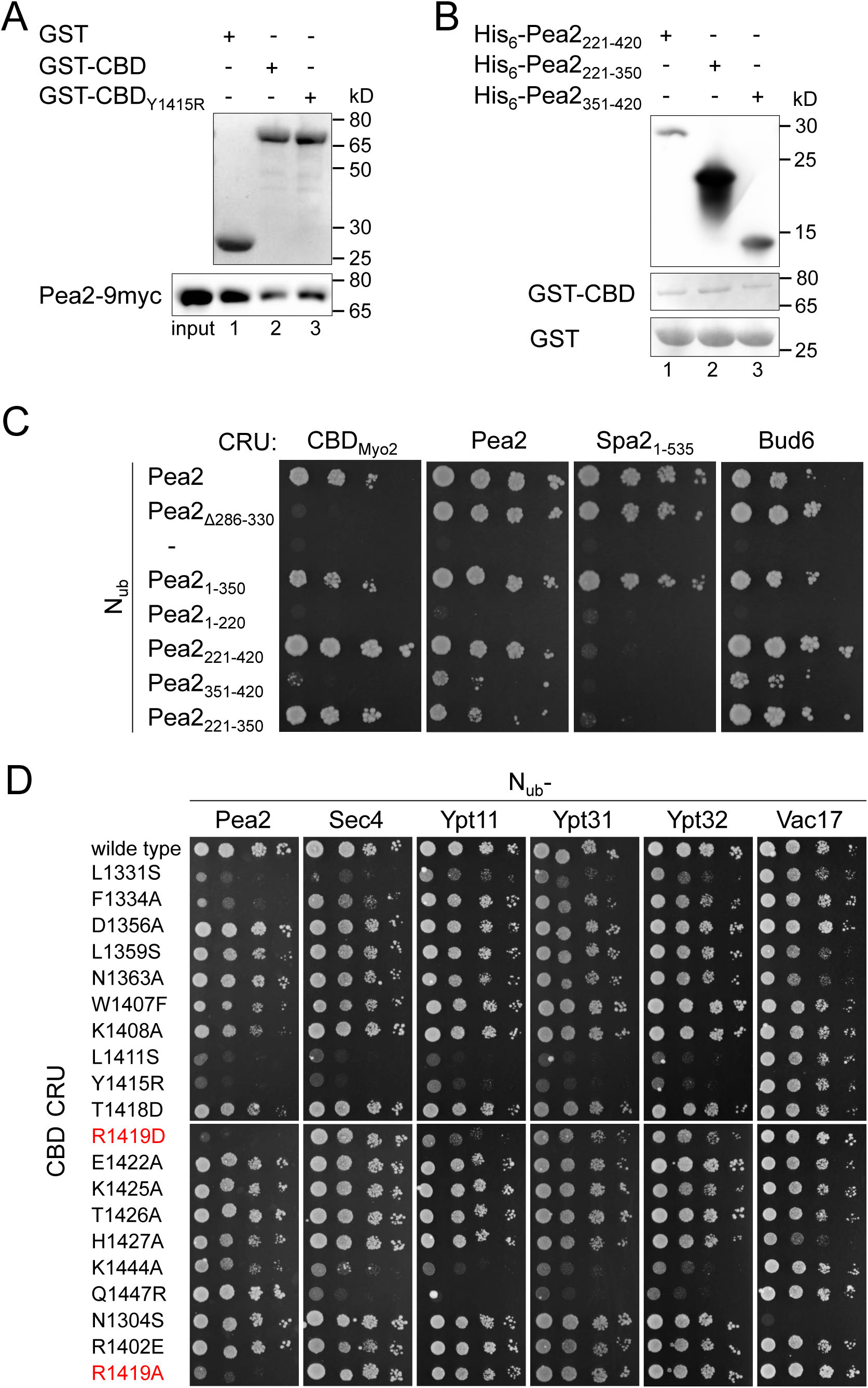
(A) Upper panel: Ponceau staining of GST (lane1) or GST-fusions to CBD (lane 2) or CBD_Y1415R_ (lane 3) used for pull downs shown in Figure 1E. Equal fractions of the elutions from Glutathion Sepharose beads were separated by SDS PAGE and transferred onto nitrocellulose. Lower panel: Anti-myc staining of unbound Pea2-9myc of the pull down shown in Figure 1E. Loading as in upper panel. (B) Upper panel: Anti-His western blot of the unbound 6His-Pea2_221-420_ (lane 1), 6His-Pea2_221-350_ (lane 2) and 6His-Pea2_351-420_ (lane 3) used in the pull-down of Figure 1F. Shown are the supernatants after incubation with immobilized GST-CBD. Lower panels: Ponceau-staining of GST-CBD and GST used for the pull-downs shown in Figure 1F after elution from beads, SDS-PAGE and transfer onto nitrocellulose. (C) Extended split-ub assay of Figure 1G with centromeric plasmids containing indicated N_ub_-Pea2 fragments in haploid yeast cells expressing CBD-, Pea2-, Spa2_1-535_ - and Bud6CRU. (D) Extended split-ubiquitin assay of Figure 2B with indicated CBD CRU mutants tested against N_ub_-Pea2, -Sec4, -Ypt11, -Ypt31, - Ypt32 and -Vac17.

**Figure S3.**
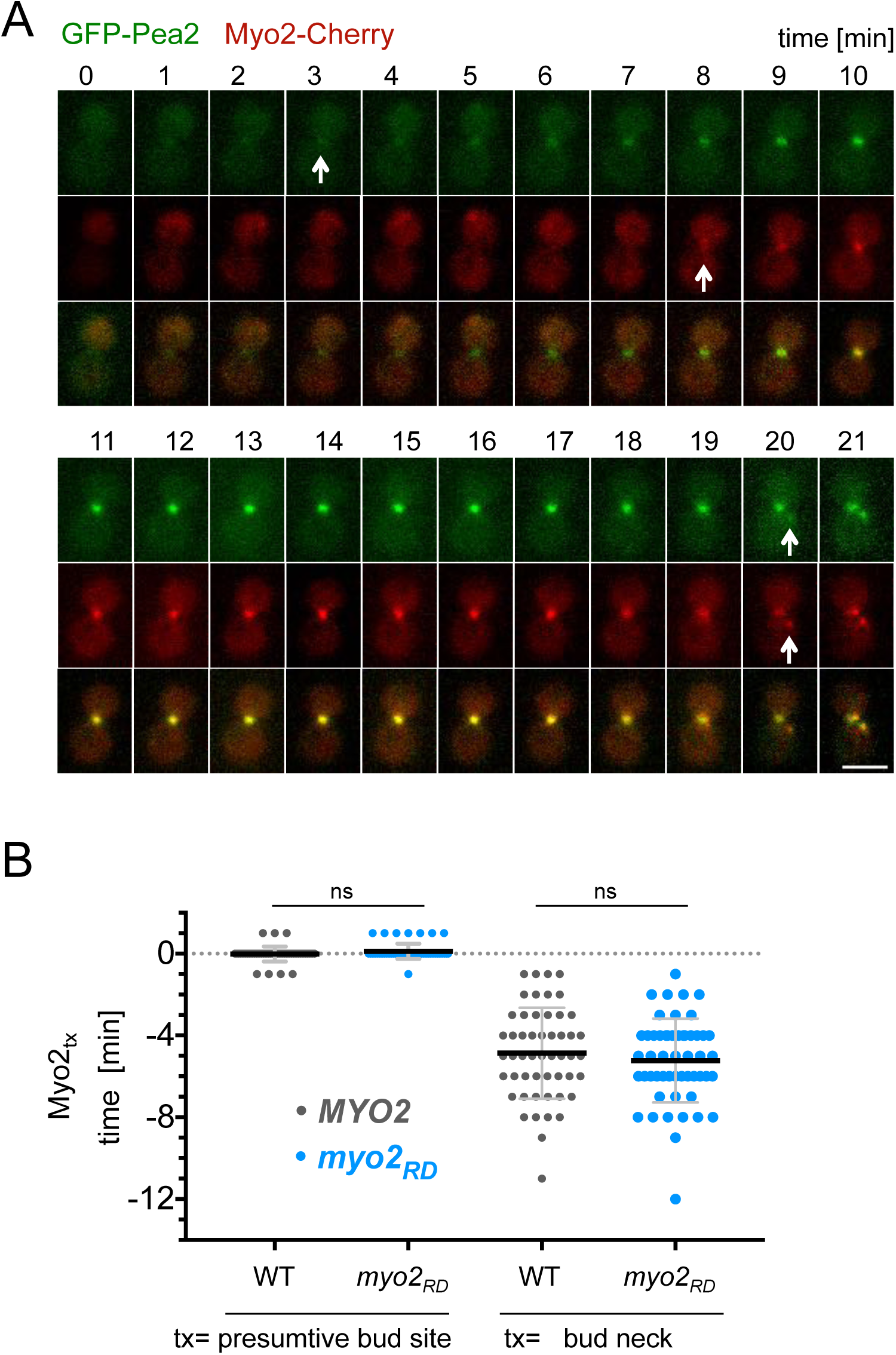
(A) Time-lapse analysis of cells co-expressing GFP-Pea2 and Myo2-mCherry in wild type cells. Arrows indicate the first visible GFP- or mCherry signal occurring during cytokinesis at the bud neck (min -5 and 0), and in G1 during assembly of the presumptive bud site (min 12). Scale bar 3 µm. (B) Appearance of Myo2-mCherry (Myo2_tx_) compared to GFP-Pea2 at the presumptive bud site and bud neck in wild type and *myo2*_*RD*_ cells. Myo2 and Pea2 appear simultaneously at the presumptive bud site in wild type- (0.02 min, n=53) and *myo2*_*RD*_-cells (−0.11 min, n=54). During cytokinesis Myo2-mCherry appears 4.87 min after GFP-Pea2 at the bud neck of wild type-cells (n=52), and 5.23 min after GFP-Pea2 at the bud neck of *myo2*_*RD*_-cells (n=53). Error bars SD.

**Figure S4.**
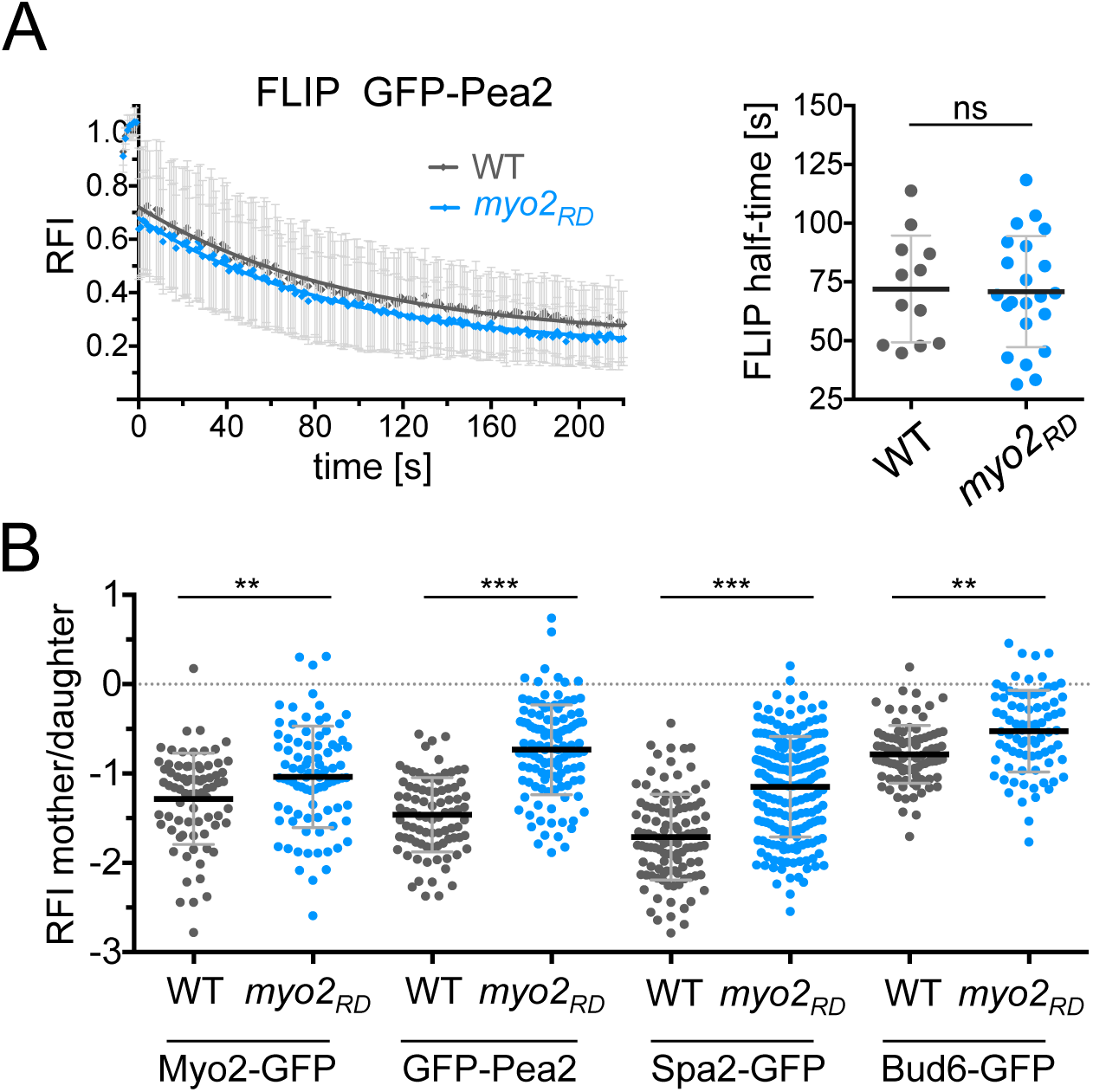
(A) FLIP measurement of GFP-Pea2 in wild type and *myo2*_*RD*_-cell. Left panel: Curves are the fitted mean of single measurements of the relative fluorescence intensity (RFI) in wild type (n=12) and *myo2*_*RD*_ (n=22) cells. Mother cells were continuously photo bleached every 5 sec and z-stack images were taken every second. Right panel: The calculated FLIP half-times of GFP-Pea2 in wild type- (72.0 sec) and *myo2*_*RD*_-cells (70.9 sec). Error bars SD. (B) Distribution of the GFP fusions to Myo2, Pea2, Spa2 and Bud6 in wild type and *myo2*_*RD*_ cells. The ratios of the RFIs of mother/daughter of Myo2-GFP (n_WT_ =75, n_*myo2RD*_ =87), GFP-Pea2 (n_WT_=87, n_*myo2RD*_=118), Spa2-GFP (n_WT_=110, n_*myo2RD*_=190) and Bud6-GFP (n_WT_=91, n_myo2RD_=80) show significant smaller values in wild type than in *myo2*_*RD*_-cells, indicating a higher concentration of the GFP fusion proteins in the bud of wild type cells.

### Supplementary Tables

**Table S1.**
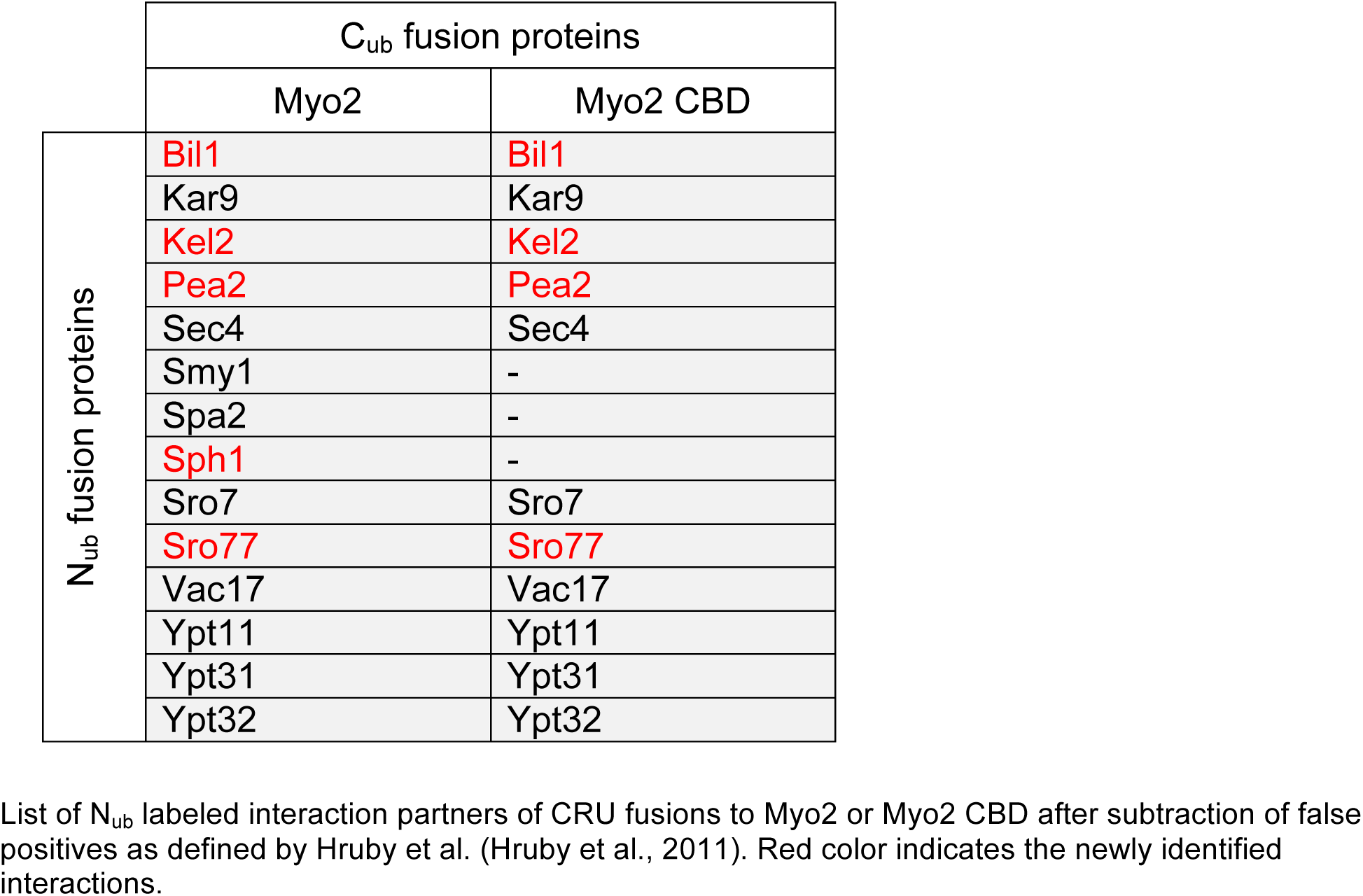

**Table S2.** Parameter values for modeling.

| parameter | value | reference |
| --- | --- | --- |
| $N$, the number of particles | 400 | this study |
| $N_{\max}$, the maximal number of cables | 10 | (Amberg, 1998; Schott et al., 2002) |
| $L_{\max}$, the maximal length of cable attachment to the cortex | 0.6 $\mu\text{m}$ | (Pruyne and Bretscher, 2000; Pruyne et al., 2004) |
| $D$, diffusion coefficient | $3 \times 10^{-5} \mu\text{m}^2/\text{s}$ | this study |
| $k_n$, cable nucleation rate | $3 \text{ s}^{-1}$ | this study |
| $k_e$, cable elongation rate | 0.3 $\mu\text{m}/\text{s}$ | (Pollard, 1986) |
| $k_d$, cable detachment rate | $0.3 \text{ s}^{-1}$ | this study |
| $k_a$, advection rate | 0.01 $\mu\text{m}/\text{s}$ | this study |

**Table S3.** List of *S. cerevisiae* strains used and created in this study.

| Strain | Name | Relevant genotype | Origin |
| --- | --- | --- | --- |
| JD47 | JD47 | MATa, *his3-Δ200, leu2-3, 112 lys2-801, trp1-Δ63, ura3-52* | Dohmen et al., 1995 |
| JD53 | JD53 | MATα, *his3-Δ200, leu2-3, 112 lys2-801, trp1-Δ63, ura3-52* | Dohmen et al., 1995 |
| JD51 | JD51 | MATa/α, *his3-Δ200/his3-Δ200, leu2-3/leu2-3, 112 lys2-801/112 lys2-801, trp1-Δ63/trp1-Δ63, ura3-52/ura3-52* | Dohmen et al., 1995 |
| ULM53 | ULM53 | MATα, *his3-Δ200, leu2-3, 112 lys2-801, trp1-Δ63, ura3-52* | Grinhagens et al., 2020 |
| YAD1685 | Myo2 CRU | JD47, *BEM1::natNT2 P<sub>MET17</sub>MYO2-C<sub>ub</sub>-R-URA3 HIS3* | This work |
| UNY177 | CBD<sub>Myo2</sub>CRU | JD47, *pP<sub>MET17</sub>MYO2<sub>1152-end</sub>-C<sub>ub</sub>-R-URA3 CEN HIS3* | This work |
| UNY178 | CBD<sub>Myo2 N1304S</sub>CRU | JD47, *pP<sub>MET17</sub>myo2<sub>N1304S_1152-end</sub>-C<sub>ub</sub>-R-URA3 CEN HIS3* | This work |
| UNY179 | CBD<sub>Myo2 R1402E</sub>CRU | *pP<sub>MET17</sub>myo2<sub>R1402E_1152-end</sub>-C<sub>ub</sub>-R-URA3 CEN HIS3* | This work |
| UNY180 | CBD<sub>Myo2 Y1415R</sub>CRU | *pP<sub>MET17</sub>myo2<sub>Y1415R_1152-end</sub>-C<sub>ub</sub>-R-URA3 CEN HIS3* | This work |
| NJY356 | Pea2 CRU | JD47, *PEA2::PEA2-C<sub>ub</sub>-R-URA3 HIS3* | Neller et al., 2015 |
| NJY355 | Bud6 CRU | JD47, *BUD6::BUD6-C<sub>ub</sub>-R-URA3 HIS3* | Glomb et al., 2019 |
| YAD397 | Spa2<sub>1-535</sub> CRU | JD47, *pP<sub>MET17</sub>SPA2<sub>1-535</sub>-C<sub>ub</sub>-R-URA3 CEN HIS3* | This work |
| YAD2608 | Myo2 | JD47, *MYO2::MYO2-hphNT1* | This work |
| YAD2611 | Myo2<sub>RA</sub> | JD47, *MYO2::myo2<sub>R1419A</sub>-hphNT1* | This work |
| YAD2614 | Myo2<sub>RD</sub> | JD47, *MYO2::myo2<sub>R1419D</sub>-hphNT1* | This work |
| YAD2647 | Myo2<sub>WT</sub> CRU | JD47, *MYO2::MYO2-C<sub>ub</sub>-R-URA3 HIS3 hphNT1* | This work |
| YAD2648 | Myo2<sub>RA</sub> CRU | JD47, *MYO2::myo2<sub>R1419A</sub>-C<sub>ub</sub>-R-URA3 HIS3 hphNT1* | This work |
| YAD2649 | Myo2<sub>RD</sub> CRU | JD47, *MYO2::myo2<sub>R1419D</sub>-C<sub>ub</sub>-R-URA3 HIS3 hphNT1* | This work |
| YAD249 | pea2Δ | JD47, *PEA2::hphNT1* | Neller et al., 2015 |
| YAD2396 | Pea2<sub>286-330Δ</sub> | JD47, *PEA2::pea2<sub>286-330Δ</sub>* | This work |
| YAD2398 | GFP-Pea2 | JD47, *P<sub>PEA2</sub>::P<sub>PEA2</sub>-GFP loxP,* | This work |
| YAD3023 | GFP-Pea2<br>Myo2-mCherry | JD47, *P<sub>PEA2</sub>::P<sub>PEA2</sub>-GFP loxP,*<br>*MYO2::MYO2-mCherry URA3 hphNT1* | This work |
| YAD2822 | GFP-Pea2 | JD47, *P<sub>PEA2</sub>::P<sub>PEA2</sub>-GFP loxP,*<br>*MYO2::MYO2-hphNT1* | This work |
| YAD2823 | GFP-Pea2<br>Myo2<sub>RD</sub> | JD47, *P<sub>PEA2</sub>::P<sub>PEA2</sub>-GFP loxP,*<br>*MYO2::myo2<sub>R1419D</sub>-hphNT1* | This work |
| YAD3025 | GFP-Pea2<br>Myo2<sub>RD</sub>-mCherry | JD47, *P<sub>PEA2</sub>::P<sub>PEA2</sub>-GFP loxP,*<br>*MYO2::myo2<sub>R1419D</sub>-mCherry URA3 hphNT1* | This work |
| YAD2654 | Myo2-GFP | JD47, *MYO2::MYO2-GFP TRP1 hphNT1* | This work |
| YAD2655 | Myo2<sub>RA</sub>-GFP | JD47, *MYO2::myo2<sub>R1419A</sub>-GFP TRP1* | This work |
|  |  | *hphNT1* |  |
| YAD2656 | Myo2<sub>RD</sub>-GFP | JD47, MYO2::myo2<sub>R1419D</sub>-GFP TRP1<br>*hphNT1* | This work |
| YAD902 | Myo2-GFP<br>pea2Δ | JD47, PEA2::hphNT1,<br>MYO2::MYO2-GFP URA3 | This work |
| YAD2657 | Spa2-GFP | JD47, SPA2::SPA2-GFP TRP1,<br>MYO2::MYO2-hphNT1 | This work |
| YAD2659 | Spa2-GFP<br>Myo2<sub>RD</sub> | JD47, SPA2::SPA2-GFP TRP1,<br>MYO2::myo2<sub>R1419D</sub>-hphNT1 | This work |
| STY1329 | Spa2-GFP<br>bni1Δ | JD47, SPA2::SPA2-GFP TRP1,<br>BNI1::hphNT1 | This work |
| STY1254 | Spa2-GFP<br>msb3Δ msb4Δ | JD47, SPA2::SPA2-GFP TRP1,<br>MSB3::hphNT1, MSB4::kanMX | This work |
| YAD3656 | Spa2-mCherry | ULM53, SPA2::SPA2-mCherry URA3,<br>MYO2::MYO2-hphNT1 | This work |
| YAD3658 | Spa2-mCherry<br>Myo2<sub>RD</sub> | ULM53, SPA2::SPA2-mCherry URA3,<br>MYO2::myo2<sub>R1419D</sub>-hphNT1 | This work |
| YAD2660 | Bud6-GFP | JD47, BUD6::BUD6-GFP TRP1,<br>MYO2::MYO2-hphNT1 | This work |
| YAD2662 | Bud6-GFP<br>Myo2<sub>RD</sub> | JD47, BUD6::BUD6-GFP TRP1,<br>MYO2::myo2<sub>R1419D</sub>-hphNT1 | This work |
| YAD3560 | GFP-Pea2<br>Spa2-mCherry | JD47, P<sub>PEA2</sub>::P<sub>PEA2</sub>-GFP loxP,<br>SPA2::SPA2-mCherry URA3 hphNT1 | This work |
| YAD3270 | GFP-Pea2<br>Myo2-mCherry | JD47, P<sub>PEA2</sub>::P<sub>PEA2</sub>-GFP loxP,<br>MYO2::MYO2-mCherry URA3 hphNT1 | This work |
| YAD3272 | GFP-Pea2<br>Myo2<sub>RD</sub>-mCherry | JD47, P<sub>PEA2</sub>::P<sub>PEA2</sub>-GFP loxP,<br>MYO2::myo2<sub>R1419D</sub>-mCherry URA3 hphNT1 | This work |
| YAD3561 | GFP-Pea2<br>Gic2<sub>PBD</sub>-tdTomato | JD47, P<sub>PEA2</sub>::P<sub>PEA2</sub>-GFP loxP,<br>ura3-52::URA3 GIC2<sub>1-208</sub>-tdTomato HIS3 | This work |
| YAD2666 | GFP-Sec4 | JD47, ura3-52::URA3 P<sub>SEC4</sub>-GFP-SEC4,<br>MYO2::MYO2-hphNT1 | This work |
| YAD2668 | GFP-Sec4<br>Myo2<sub>RD</sub> | JD47, ura3-52::URA3 P<sub>SEC4</sub>-GFP-SEC4,<br>MYO2::myo2<sub>R1419D</sub>-hphNT1 | This work |
| YAD3459 | Pex-mCherry-Pea2<br>GFP-SKL | JD47, pea2::kanMX, ura3-52::URA3<br>P<sub>MET17</sub>PEX3<sub>1-45</sub>-mCherry-PEA2,<br>MYO2::MYO2-hphNT1,<br>pP<sub>CTA1</sub>-GFP-SKL LEU2 CEN | This work |
| YAD3460 | Pex-mCherry-Pea2<br>Myo2<sub>RD</sub><br>GFP-SKL | JD47, pea2::kanMX, ura3-52::URA3<br>P<sub>MET17</sub>PEX3<sub>1-45</sub>-mCherry-PEA2,<br>MYO2::myo2<sub>R1419D</sub>-hphNT1,<br>pP<sub>CTA1</sub>-GFP-SKL LEU2 CEN | This work |
| YAD3461 | Pex-mCherry<br>GFP-SKL | JD47, pea2::kanMX, ura3-52::URA3<br>P<sub>MET17</sub>PEX3<sub>1-45</sub>-mCherry,<br>MYO2::MYO2-hphNT1,<br>pP<sub>CTA1</sub>-GFP-SKL LEU2 CEN | This work |
| YAD3462 | Pex-mCherry<br>Myo2<sub>RD</sub><br>GFP-SKL | JD47, pea2::kanMX, ura3-52::URA3<br>P<sub>MET17</sub>PEX3<sub>1-45</sub>-mCherry,<br>MYO2::myo2<sub>R1419D</sub>-hphNT1,<br>pP<sub>CTA1</sub>-GFP-SKL LEU2 CEN | This work |
| YAD941 | Pea2-9myc | JD47, PEA2::hphNT1,<br>pP<sub>MET17</sub> PEA2-9xMYC 315 | This work |

**Table S4.**
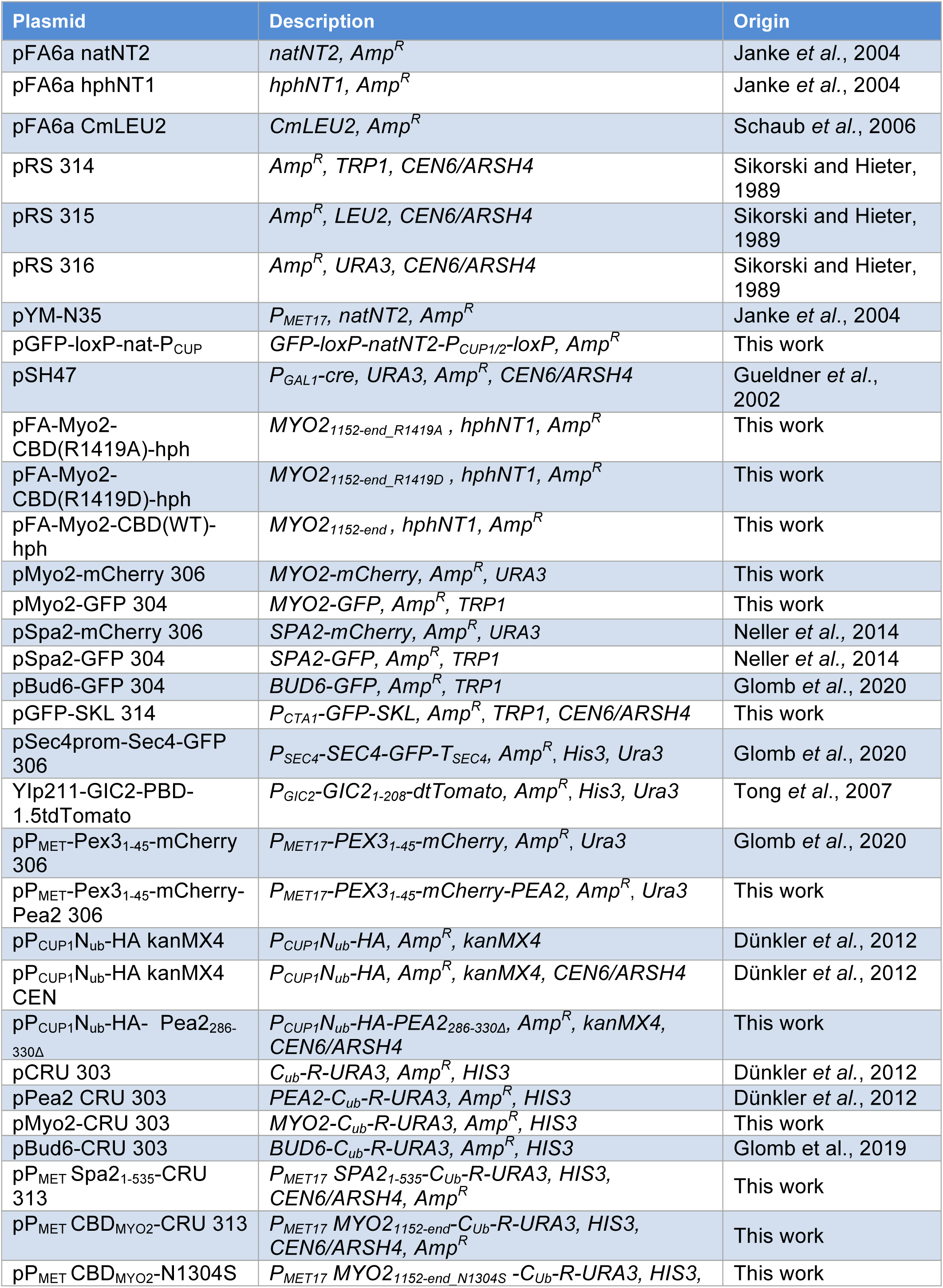
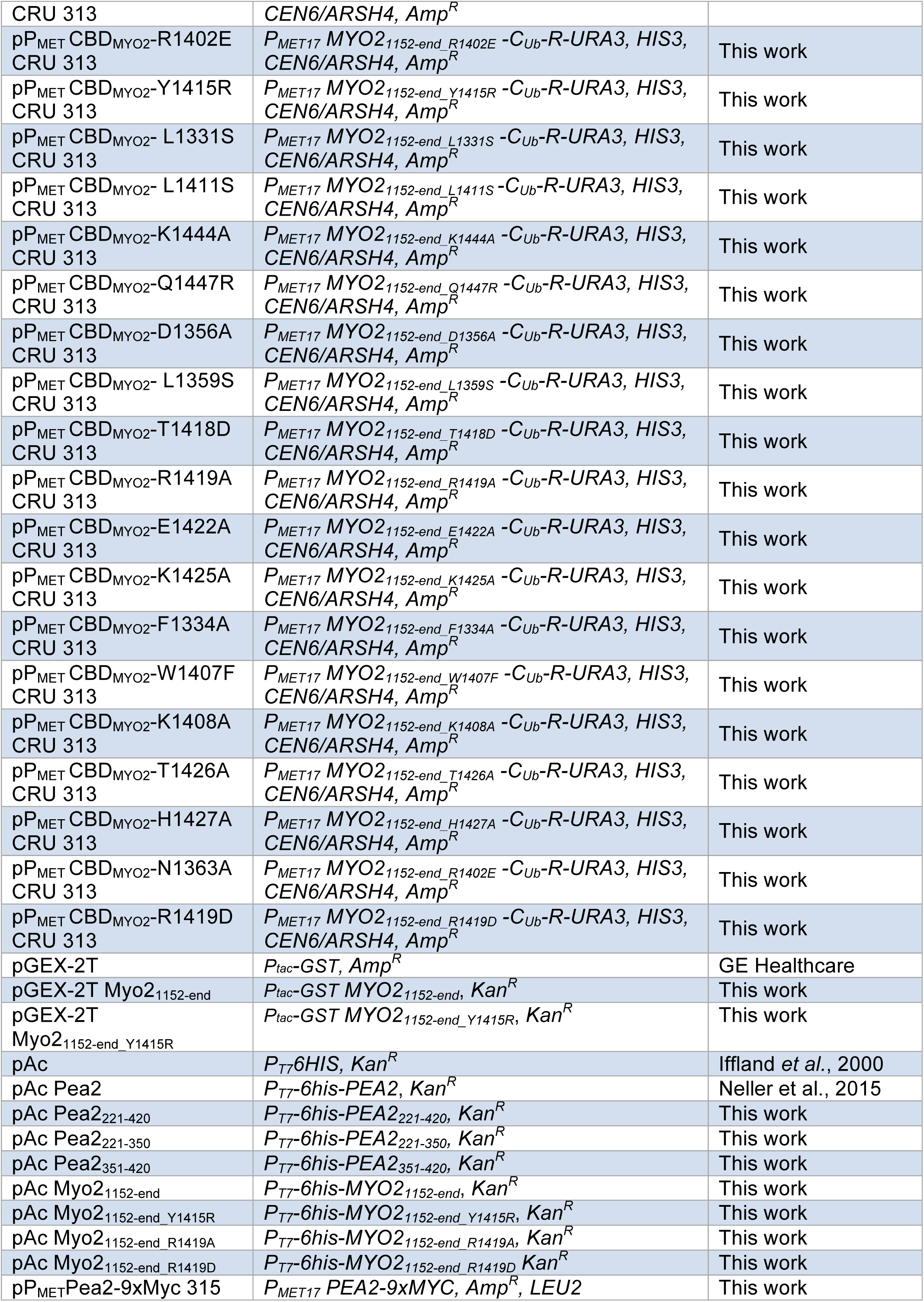
List of constructed plasmids in this study.

### Movies

**Movie S1**

Time-lapse analysis of yeast cells expressing GFP-Pea2 and Myo2-mCherry. Frames were taken every 2 min and cells incubated at 30°C. Scale bar 5 µm.

**Movie S2**

Time-lapse analysis of yeast cells expressing GFP-Pea2 and Myo2_RD_-mCherry. Frames were taken every 2 min and cells incubated at 30°C. Scale bar 5 µm.

**Movie S3**

Mating of wild type a- and α-cells expressing Spa2-GFP or Spa2-mCherry respectively at 30°C. Frames were taken every 2 min. Scale bar 5 µm.

**Movie S4**

Mating of *MATa*-myo2_RD_-cells expressing Spa2-GFP with *MATα*-wild type cells expressing Spa2-mCherry at 30°C. Frames were taken every 2 min. Scale bar 5 µm.

